# Assessing the effects of thinning on stem growth allocation of individual Scots pine trees

**DOI:** 10.1101/2020.03.02.972521

**Authors:** Ninni Saarinen, Ville Kankare, Tuomas Yrttimaa, Niko Viljanen, Eija Honkavaara, Markus Holopainen, Juha Hyyppä, Saija Huuskonen, Jari Hynynen, Mikko Vastaranta

## Abstract

Forest management alters the growing conditions and thus further development of trees. However, quantitative assessment of forest management on tree growth has been demanding as methodologies for capturing changes comprehensively in space and time have been lacking. Terrestrial laser scanning (TLS) has shown to be capable of providing three-dimensional (3D) tree stem reconstructions required for revealing differences between stem shapes and sizes. In this study, we used 3D reconstructions of tree stems from TLS and an unmanned aerial vehicle (UAV) to investigate how varying thinning treatments and the following growth effects affected stem shape and size of Scots pine (*Pinus sylvestris* L.) trees. The results showed that intensive thinning resulted in more stem volume and therefore total biomass allocation and carbon uptake compared to the moderate thinning. Relationship between tree height and diameter at breast height (i.e. slenderness) varied between both thinning intensity and type (i.e. from below and above) indicating differing response to thinning and allocation of stem growth of Scots pine trees. Furthermore, intensive thinning, especially from below, produced less variation in relative stem attributes characterizing stem shape and size. Thus, it can be concluded that thinning intensity, type, and the following growth effects have an impact on post-thinning stem shape and size of Scots pine trees. Our study presented detailed measurements on post-thinning stem growth of Scots pines that have been laborious or impracticable before the emergence of detailed 3D technologies. Moreover, the stem reconstructions from TLS and UAV provided variety of attributes characterizing stem shape and size that have not traditionally been feasible to obtain. The study demonstrated that detailed 3D technologies, such as TLS and UAV, provide information that can be used to generate new knowledge for supporting forest management and silviculture as well as improving ecological understanding of boreal forests.

## Introduction

Allocation of photosynthesis products between different parts of a tree define their development. If resources are limited, trees first channel them towards respiration and sustaining existing parts, producing fine roots and seed (i.e. reproduction), height growth and only then growth in diameter at breast height (1.3 m, DBH). Competition for resources (e.g. light, water, nutrients) between trees also affects development of different tree parts. Thus, differences in the allocation of photosynthesis products within a tree and competition between trees result in variation in tree architecture.

In thinning, as a part of forest management, competition within a population is regulated as part of the trees are removed. Therefore, ecologically thinning is aimed at improving growing conditions (i.e., light, temperature, water, nutrients) of remaining trees and economically to maximize the net present value of a stand by decreasing the opportunity costs of the capital. Especially when part of shadowing leaf mass is removed in thinning, the amount of light is increased, and growth of remaining trees is enhanced (White 1980). Additionally, an increase in the amount of nutrients, especially in nitrogen, have been demonstrated after thinning in Scots pine (*Pinus sylvestris* L.) stands due to logging residue (Kimmins & Scoullar 1983, Kukkola & Mälkönen 1997). The development and profitability of forestry can be affected through thinning intensity, type (i.e. selection of the remaining and removing trees) as well as timing. More intensive thinning signifies fewer remaining trees and decrease in standing volume (Mäkinen & Isomäki 2004a).

Intermediate and suppressed trees can expect to benefit from thinning more than co-dominant and dominant trees as growing conditions are improved and competition decreased (i.e. more available light) for trees in lower canopy layers. Mäkinen & Isomäki (2004c) reported that relative growth of basal area after thinning was independent of tree size, but it was the largest among intermediate and suppressed trees compared to co-dominant and dominant trees. Conversely, height growth of trees in forests with low tree density is typically smaller than in forests with normal tree density. Especially in Scots pine stands, growth of dominant height can temporarily decline (during 25 years on average) after heavy thinning from below (Mäkinen & Isomäki 2004a). Thinning affects the forest structure and relationship between the remaining trees. For example, intensive thinning homogenized DBH distributions in Scots pine stands as dense forests had larger variation in DBH (Mäkinen & Isomäki (2004c). Similarly, it has been shown that DBH growth is affected by tree density and the available space, for example, according to Savill & Sandels (1983) more spacing (i.e. less trees per ha) increased DBH growth.

Studies on diameter growth along the stem of conifers reveal that width of annual rings in the most top part of a stem is relatively narrow but increases linearly with leaf volume (i.e. lower stem part) (Farrar 1961, Kozlowski 1971). In these studies, it has been shown that in dominant and open-grown trees the annual ring thickens near the bottom of a stem whereas in suppressed trees the ring throughout the stem is thinner than in dominant trees. Farrar (1961) reported increasing annual ring growth in the bottom part of a stem after thinning while growth in top part of a stem (i.e. within crown) was comparable to the situation before thinning. Similarly, Valinger (1992) demonstrated that thinning promoted diameter growth in the bottom part of a stem and decreased height growth. Mäkinen & Isomäki (2004b), on the other hand, demonstrated that diameter growth of Norway spruce (*Picea abies* (L.) H. Karst) concentrated on top part of a stem in dense forests whereas in forests where trees had more space available, diameter growth was similar in different heights along a stem.

When impacts of thinning intensity were assessed through attributes characterizing stem shape of Scots pine trees, Mäkinen & Isomäki (2004c) reported that height/diameter ratio (i.e. slenderness) decreased and stem taper (i.e. DBH – diameter at 6 m height) increased when thinning intensity increased from unthinned to light, moderate, and heavy thinning (i.e. average basal area after the thinning treatment compared to the state before thinning ≥95%, 80-94%, 65-79%, and ≤64% of control plots, respectively). Similarly, crown height and crown ratio were significantly different between thinning intensities (Mäkinen & Isomäki 2004c). It should be noted that the thinning intensities, even the heavy thinning, studied in Mäkinen & Isomäki (2004c) correspond approximately thinning intensities applied nowadays in Finland (Rantala 2011).

In addition to different thinning intensities, thinning type defines what kind of trees are removed and most importantly left to grow. In thinning from below, intermediate and suppressed as well as overgrown trees are removed in order to increase the volume growth of the remaining trees whereas in thinning from above dominant and overgrown trees are removed. At stand level, thinning from above increased basal area and volume growth in Scots pine stands compared to thinning from below whereas thinning from below increased growth of dominant height more than thinning from above (Vuokila 1977). Similarly, Mielikäinen & Valkonen (1991) reported that thinning from above resulted in 4-8% more volume growth in Scot pine stands than thinning from below. Furthermore, Eriksson & Karlsson (1997) illustrated that thinning from above did not decrease volume growth in Scots pine stands in comparison with thinning from below.

Growing conditions are, thus, dependent on tree density that affects growth and structure of trees (Harper 1977) whereas changes in height and diameter growth alter stem shape. Therefore, understanding changes in stem shape enable assessing the effects of thinning on future growth as well as possible susceptibility of snow and wind damage of remaining trees. However, studies reviewed above were only based on a limited number of observations and the observations used were only able to capture limited changes along a stem. Observations only included DBH, diameter at 6-m height, and tree height measured with calipers and clinometers. Differences in stem volume were predicted using allometric equations that are incapable of considering variation in stem shape. Thus, our understanding of the effects of forest management on details of stem shape and growth responses is still rather limited.

Terrestrial laser scanning (TLS) has proven to non-destructively provide three-dimensional (3D) information on tree stems (Liang et al. 2014, Kankare et al. 2013, Raumonen et al. 2013, Saarinen et al. 2017) that has not been possible to obtain with calipers or measurement tape. Individual trees can be detected from a TLS-based point cloud through identification of circular shapes (Aschoff et al. 2004, Maas et al. 2008) or clusters of points (Cabo et al. 2018, Zhang et al. 2019). Points from individual trees can then be utilized in reconstructing the entire architectural structure of a tree (Raumonen et al. 2013, Hackenberg et al. 2014) or only the stem (Liang et al. 2011, Heinzel & Huber 2017). When acquisition and preprocessing of TLS data are thoroughly carried out, millimeter-level details can be observed from TLS point clouds (Wilkes et al. 2017, Liang et al. 2018), which enables accurate geometrical tree reconstruction (Hackenberg et al. 2014). In applications related to boreal forests, TLS has been utilized in quantifying stem growth and changes in stem taper (Luoma et al. 2019), reparametrizing an existing taper curve model (Saarinen et al. 2019), and assessing timber quality (Pyörälä et al. 2019). In Central Europe, on the other hand, impacts of silviculture on stem shape of Norway spruce (Jacobs et al. 2019) and European beech (Juchheim et al. 2017b, Georgi et al. 2018, Noyer et al. 2019) have been studied. In Juchheim et al. (2017b) and Georgi et al. (2018), several attributes characterizing stem size and shape were generated from TLS point clouds (e.g. tree height, DBH, stem volume, crown-base height, height-diameter ratio, taper, lean, and sweep) of which most are attributes that have been generated also from traditional measurements. However, as shown by Jacobs et al. (2019) and Saarinen et al. (2019), taper curve is possible to obtain from TLS point clouds and it enables generation of unconventional attributes that may be used in revealing differences in the stem development as well as stem shape and size.

It has been, however, reported that reliable estimates in tree height have been challenging to obtain in boreal forests (Liang et al. 2018) even when scan positions are selected for the best possible visibility of treetops (Saarinen et al. 2017). Photogrammetric point clouds from unmanned aerial vehicles (UAVs), on the other hand, have proven to be an attractive option for characterizing forest structure (Puliti et al. 2015, Wallace et al. 2016, Goodbody et al. 2017, Alonzo et al. 2018, Saarinen et al. 2018, Kotivuori et al. 2020). Measurement geometry of UAVs enable capturing of the upper canopy and combination of point clouds from TLS and UAV could further enhance characterization of forest structure as well as individual trees.

Although theoretical understanding exists on the effects of forest management and resulting growing conditions on allocation between height and diameter growth of trees, understanding differences in tree architecture based on quantitative information is still limited. However, detailed information on stem shape and size has not been available for studies investigating effects of both thinning intensity and type. Therefore, the objective of this study is to investigate the effects of intensity and type of thinning on post-thinning stem growth of Scots pine trees. Point clouds from TLS and UAV provid an opportunity in generating various new attributes characterizing both absolute and relative stem shape and size. Based on previous studies on growth and yield, we hypothesized that thinning intensity and type result in differing stem shape and size. It was further divided to following research questions: **(1)** how thinning intensity and type affect stem shape and size; **(2)** how thinning intensity and type affect growth allocation between DBH and tree height; and **(3)** how thinning intensity and type affect variation in stem shape and size.

## Materials and methods

### Study site and data acquisition

The research was conducted in southern Finland in three study sites dominated by Scots pine (Palomäki, Pollari, and Vesijako) (Figure 1) and established and maintained by Natural Resources Institute Finland (Luke). Study site in Palomäki was established in 2005 whereas study sites in Pollari and Vesijako in 2006. The study sites are located in the same vegetation zone namely southern Boreal forest zone at relative flat terrain, the elevation is 135 m, 155 m, and 120 m above sea level in Palomäki, Pollari, and Vesijako, respectively. Furthermore, the temperature sum is 1195, 1130, and 1256 degree days in Palomäki, Pollari, and Vesijako, respectively. Each study site is characterized as mesic heath forest (i.e., Myrtillus forest site type according to Cajander (1909)) and includes nine rectangular sample plots with size varying from 1000 m^2^ to 1200 m^2^ resulting in total of 27 sample plots. At the time of establishment, the stand age was 50, 45, and 59 years for Palomäki, Pollari, and Vesijako, respectively. Also, first thinning had been carried out in early 1990s for all study sites.

**Figure 1.**
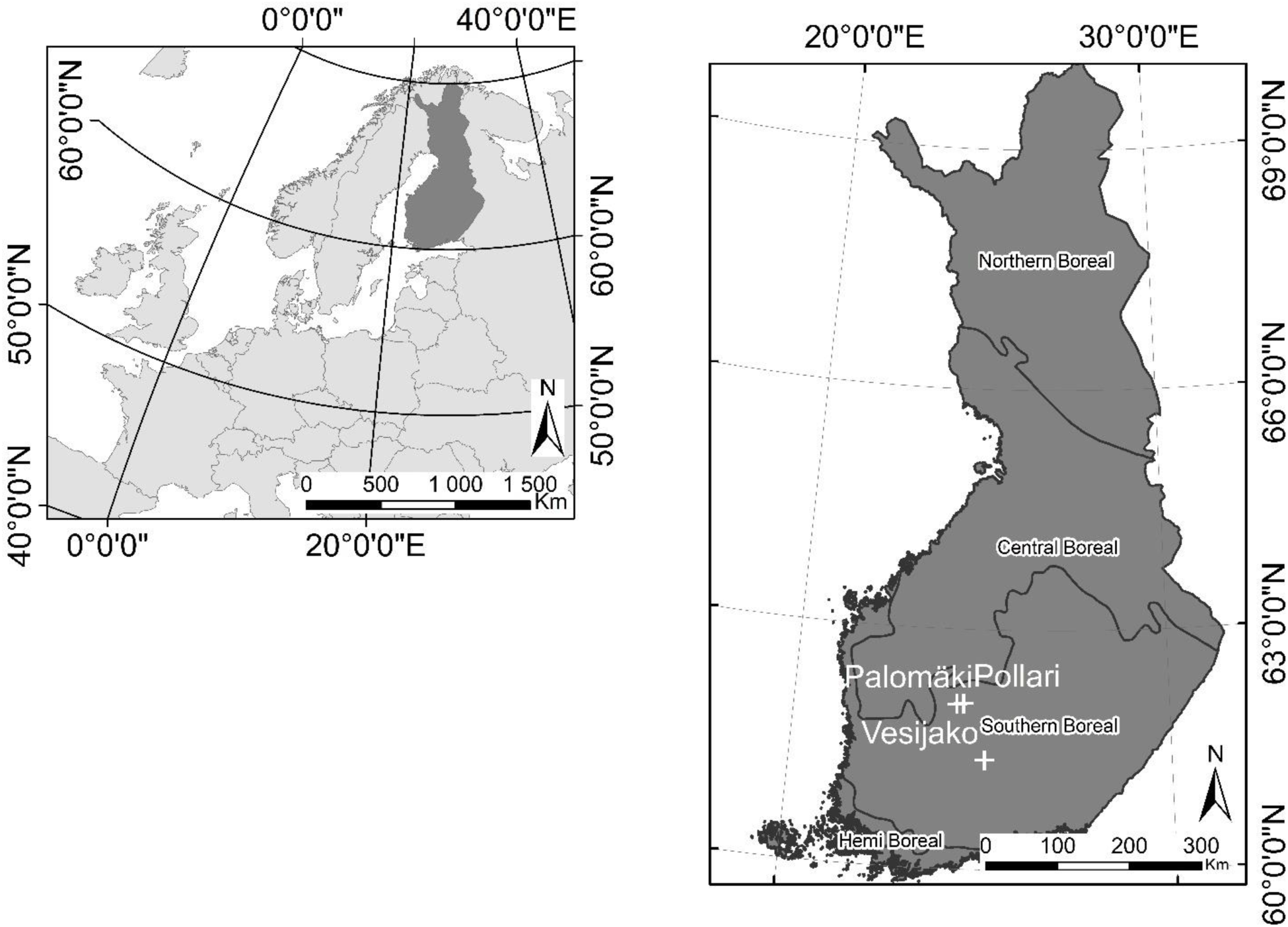
Location of the three study sites namely Palomäki, Pollari, and Vesijako and vegetation zones in Finland.

The experimental design of the study sites includes two levels of thinning intensity and three thinning types resulting in six different thinning treatments, namely i) moderate thinning from below, ii) moderate thinning from above, iii) moderate systematic thinning from above, iv) intensive thinning from below, v) intensive thinning from above, and vi) intensive systematic thinning from above, as well as a control plot where no thinning has been carried out since the establishment. Moderate thinning refers to prevailing thinning guidelines applied in Finland (Rantala 2011) whereas intensive thinning corresponds 50% lower remaining basal area (m^2^/ha) than in the plots with moderate thinning intensity. Regarding thinning types, small and suppressed trees as well as unsound and damaged trees (e.g., crooked, forked) were first removed from plots where thinning from below or above was carried out. Furthermore, suppressed and co-dominant trees were removed in thinning from below whereas mostly dominant trees were removed in thinning from above, but also maintaining regular spatial distribution of trees. In systematic thinning from above, on the other hand, only dominant trees were removed and small, suppressed trees were left to grow and regularity of spatial distribution of remaining trees was not emphasized similarly to other thinning types, although large gaps were avoided.

When the thinning trials were established, the plots were measured and thinning treatments were performed. The study sites have been re-measured three times after the establishment, the latest measurements were carried out in October 2018 in study site Pollari whereas study sites Palomäki and Vesijako were measured in April 2019. Thus, the time after the thinning treatments and the period for growth response on the remaining trees was 13 years in Palomäki and 12 in Pollari and Vesijako. Tree species, DBH from two perpendicular directions, crown layer, and health status were recorded from each tree within a plot for each measurement time. The proportion of Norway spruce and deciduous trees (i.e. *Betula* sp and *Alnus* sp) from the total stem volume of all trees within the 27 sample plots was 3.06% and 0.03%, respectively. Each sample plot also includes 22 sample trees, on average, from which tree height, crown base height, and height of the lowest dead branch were also measured. Stand attributes before and after thinning treatments together with thinning removals are presented in Table 1 and the development of tree-level attributes for each thinning treatment is found in Table 2. The remaining relative stand basal area after moderate thinning was ∼68% of the stocking before thinning and intensive thinning reduced stocking levels down to 34%. There were no large differences in remaining basal area or volume between thinning types with the same thinning intensity (Table 1).

**Table 1.** Mean stand characteristics by treatments before and after thinning as well as thinning removal. N = stem number per hectare, G = basal area, D_w_ = mean diameter weighted by basal area, H_w_ = mean height weighted by basal area, H_100_ = dominant height, and V = volume.

| Before thinning (2005-2006) |  |  |  |  |  |  |  |
| --- | --- | --- | --- | --- | --- | --- | --- |
|  | Moderate<br>below | Moderate<br>above | Moderate<br>systematic | Intensive<br>below | Intensive<br>above | Intensive<br>systematic | No<br>treatment |
| N/ha | 1269 | 1390 | 1348 | 1244 | 1161 | 1220 | 1337 |
| G (m<sup>2</sup>/ha) | 26.5 | 27.3 | 25.4 | 26.5 | 23.9 | 26.0 | 27.7 |
| D<sub>w</sub> (cm) | 17.5 | 17.3 | 17.1 | 18.0 | 17.6 | 18.0 | 17.8 |
| H<sub>w</sub> (m) | 16.1 | 15.9 | 15.7 | 16.3 | 15.6 | 16.2 | 16.1 |
| H<sub>100</sub> (m) | 17.3 | 17.2 | 17.1 | 17.7 | 17.0 | 17.5 | 17.5 |
| V (m<sup>3</sup>/ha) | 213.4 | 213.5 | 197.6 | 215.7 | 185.6 | 210.9 | 224.0 |
| After thinning (2005-2006) |  |  |  |  |  |  |  |
|  | Moderate<br>below | Moderate<br>above | Moderate<br>systematic | Intensive<br>below | Intensive<br>above | Intensive<br>systematic | No<br>treatment |
| N/ha | 716 | 937 | 1033 | 289 | 461 | 520 | 1337 |
| G (m<sup>2</sup>/ha) | 18.1 | 18.2 | 17.9 | 8.7 | 8.7 | 8.6 | 27.7 |
| D<sub>w</sub> (cm) | 18.7 | 16.9 | 16.1 | 20.4 | 16.5 | 15.7 | 17.8 |
| H<sub>w</sub> (m) | 16.5 | 15.8 | 15.4 | 16.9 | 15.4 | 15.5 | 16.1 |
| H<sub>100</sub> (m) | 17.3 | 16.7 | 16.6 | 17.5 | 16.2 | 16.3 | 17.5 |
| V (m<sup>3</sup>/ha) | 148.3 | 142.3 | 137.1 | 72.9 | 67.1 | 67.6 | 224.0 |

| Thinning removal (2005-2006) |  |  |  |  |  |  |  |
| --- | --- | --- | --- | --- | --- | --- | --- |
|  | Moderate<br>below | Moderate<br>above | Moderate<br>systematic | Intensive<br>below | Intensive<br>above | Intensive<br>systematic | No<br>treatment |
| N/ha | 554 | 453 | 315 | 955 | 700 | 700 | 0.0 |
| G (m<sup>2</sup>/ha) | 8.4 | 9.1 | 7.5 | 17.8 | 15.1 | 17.4 | 0.0 |
| D<sub>w</sub> (cm) | 14.7 | 18.2 | 19.7 | 16.8 | 18.3 | 19.1 | 0.0 |
| H<sub>w</sub> (m) | 15.1 | 16.1 | 16.5 | 15.9 | 15.8 | 16.5 | 0.0 |
| H<sub>100</sub> (m) | 16.1 | 17.0 | 17.0 | 17.3 | 17.0 | 17.5 | 0.0 |
| V (m<sup>3</sup>/ha) | 65.1 | 71.2 | 60.6 | 142.9 | 118.5 | 143.2 | 0.0 |

**Table 2.** Minimum, mean, maximum, and standard deviation (std) of tree-level attributes for each treatment at the year of establishment (First measurement) and last measurement. DBH = diameter at breast height.

| First measurement (2005-2006) |  |  |  |  |  |  |  |  |
| --- | --- | --- | --- | --- | --- | --- | --- | --- |
|  |  | Moderate<br>below | Moderate<br>above | Moderate<br>systematic | Intensive<br>below | Intensive<br>above | Intensive<br>systematic | No<br>treatment |
| DBH (cm) | min | 10.3 | 8.7 | 5.9 | 11.7 | 9.3 | 7.2 | 5.8 |
|  | mean | 17.6 | 15.3 | 14.8 | 19.3 | 15.1 | 14.8 | 15.4 |
|  | max | 28.2 | 28.6 | 29.5 | 29.8 | 27.2 | 28.2 | 30.7 |
|  | std | 3.3 | 3.3 | 3.5 | 3.4 | 3.0 | 4.1 | 4.6 |
| Height (m) | min | 11.9 | 12.5 | 8.0 | 13.0 | 11.3 | 10.0 | 9.7 |
|  | mean | 15.9 | 15.3 | 14.6 | 16.5 | 14.8 | 14.6 | 14.7 |
|  | max | 19.9 | 19.3 | 22.3 | 20.5 | 19.1 | 19.9 | 23.0 |
|  | std | 1.9 | 1.2 | 1.9 | 1.8 | 1.8 | 2.6 | 2.6 |
| Volume (dm<sup>3</sup>) | min | 50.9 | 39.4 | 12.7 | 75.6 | 40.2 | 23.0 | 13.0 |
|  | mean | 202.7 | 149.6 | 138.1 | 249.1 | 141.8 | 145.6 | 160.5 |
|  | max | 549.4 | 583.8 | 689.8 | 623.0 | 521.7 | 549.4 | 783.6 |
|  | std | 89.3 | 76.2 | 77.8 | 107.0 | 73.4 | 97.1 | 119.7 |
| Last measurement (2018-2019) |  |  |  |  |  |  |  |  |
|  |  | Moderate<br>below | Moderate<br>above | Moderate<br>systematic | Intensive<br>below | Intensive<br>above | Intensive<br>systematic | No<br>treatment |
| DBH (cm) | min | 13.4 | 9.1 | 7.6 | 17.9 | 13.1 | 10.7 | 6.5 |
|  | mean | 22.2 | 19.3 | 18.8 | 26.4 | 21.1 | 20.8 | 18.7 |
|  | max | 35.3 | 32.3 | 33.9 | 36.4 | 31.1 | 35.3 | 34.4 |
|  | std | 3.7 | 4.3 | 4.2 | 3.9 | 3.5 | 4.3 | 5.0 |
| Height (m) | min | 14.7 | 15.6 | 8.4 | 18.2 | 14.9 | 13.6 | 13.2 |
|  | mean | 21.2 | 20.4 | 19.4 | 21.2 | 19.1 | 19.6 | 20.0 |
|  | max | 25.9 | 25.2 | 26.0 | 25.3 | 24.1 | 25.9 | 30.3 |
|  | std | 2.1 | 1.6 | 2.2 | 1.7 | 1.5 | 2.8 | 3.0 |
| Volume (dm<sup>3</sup>) | min | 116.6 | 51.0 | 21.3 | 231.9 | 103.9 | 61.6 | 20.9 |
|  | mean | 408.3 | 306.4 | 282.5 | 563.8 | 335.2 | 347.0 | 299.4 |
|  | max | 1050.8 | 923.3 | 1039.5 | 1146.2 | 815.6 | 1050.8 | 1266.4 |
|  | std | 160.3 | 145.6 | 137.2 | 202.5 | 125.4 | 173.3 | 190.8 |

TLS data acquisition was carried out with Trimble TX5 3D laser scanner (Trible Navigation Limited, USA) for all three study sites between September and October 2018. Eight scans were place to each sample plot, the scan setup is depicted in Figure 2, and scan resolution of point distance approximately 6.3 mm at 10-m distance was used. Artificial constant sized spheres (i.e. diameter of 198 mm) were placed around sample plots and used as reference objects for registering the eight scans onto a single, aligned coordinate system. The registration was carried out with FARO Scene software (version 2018) with a mean distance error of 2.9 mm and standard deviation 1.2 mm, mean horizontal error was 1.3 mm (standard deviation 0.4 mm) and mean vertical error 2.3 mm (standard deviation 1.2 mm).

**Figure 2.**
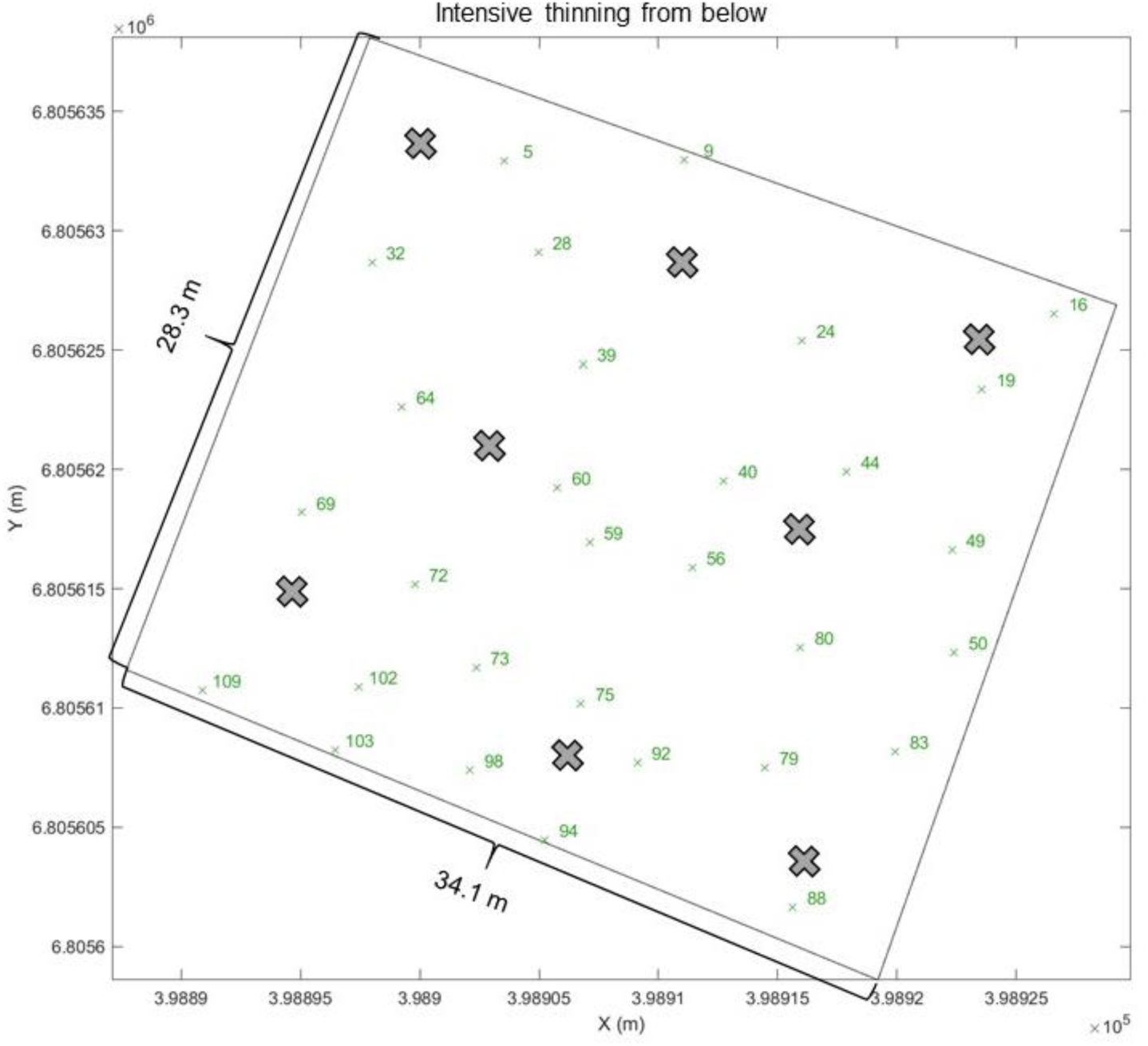
Scan design of eight scans (denoted as x) per an example sample plot thinned intensively from below. X and y axes present the coordinates of the sample plot in EUREF-FIN.

In addition to TLS data, aerial images were obtained by using an UAV with Gryphon Dynamics quadcopter frame. Two Sony A7R II digital cameras that had CMOS sensor of 42.4 MP, with a Sony FE 35mm f/2.8 ZA Carl Zeiss Sonnar T* lens, were mounted on the UAV in +15° and -15° angles. Images was acquired in every two seconds and image locations were recorded for each image. The flights were carried out on October 2, 2018. For each study site, eight ground control points (GCPs) were placed and measured. Flying height of 140 m and a flying speed of 5 m/s were selected for all the flights, resulting in 1.6 cm ground sampling distance. Total of 639, 614, and 663 images were captured for study site Palomäki, Pollari, and Vesijako, respectively, resulting in 93% and 75% forward and side overlaps, respectively. Photogrammetric processing of aerial images was carried out following the workflow as presented in Viljanen et al. (2018). The processing produced photogrammetric point clouds for each study site with point density of 804 points/m^2^, 976 points/m^2^, and 1030 points/m^2^ for Palomäki, Pollari, and Vesijako, respectively.

## Methods

The point clouds from TLS and UAV were combined for characterizing tree height and crown as comprehensively as possible. Individual tree detection and characterization from the combined point clouds followed the methodology presented by Yrttimaa et al. (2019, 2020. The process included four steps: 1) point cloud normalization, 2) tree segmentation, 3) point cloud classification, and 4) generation of tree-level attributes. The TLS point clouds were normalized (i.e. point heights were transformed to heights above ground) (1) using the *lasground* tool in LAStools software (Isenburg 2019) whereas RGB point clouds from the UAV were normalized with digital terrain model (DTM) that was based on airborne laser scanning data and provided by National Land Survey of Finland. Canopy height models (CHMs) at a 20-cm resolution were generated from the normalized UAV point clouds to segment individual tree crowns (2). Variable Window Filter approach (Popescu & Wynne 2004) was used to identify the tree top positions, and Marker-Controlled Watershed Segmentation (Meyer & Beucher 1990) was applied to delineate the tree crown segments from the CHMs. The TLS point clouds were then split into tree-segments according to the extracted UAV crown segments using point-in-polygon approach. The tree-segmented TLS point clouds were then classified (3) into stem points and non-stem points. Points that represented planar, vertical, and cylindrical surfaces were classified as stem points while the rest of the points were classified as non-stem points. Finally, tree-level attributes such as DBH, height, stem volume, and taper curve were generated (4) for each tree. Tree height was measured from the UAV point cloud while taper curve was based on diameter measurements along the stem points by fitting circles on horizontal stem point slices in every 20 cm. Initial diameter estimates were filtered by omitting clear outliers utilizing an iterative approach where diameters differing more than three median absolute deviation from the median diameter were omitted. A cubic spline curve was used to interpolate the missing diameters (especially within a crown) and to generate the final taper curve. Tree height from the UAV was utilized when fitting the cubic spline function. DBH was measured from the taper curve at the height of 1.3 m while stem volume was estimated using Huber formula by considering the stem as a sequence of vertical cylinders at 10 cm intervals.

Difference in stem shape and size was assessed between thinning treatments using tree attributes derived from information provided by the taper curves. Used attributes and their descriptions are presented in Table 3. Form factor at breast height (ff), tapering (taper), and slenderness (slend) have been used in assessing stem shape and were also included in this study. Cumulative volume (e.g. height at which 50% of stem volume accumulated (h_vol50), and volume percentiles (p10, .., p90)) enable more detailed quantitative assessment of stem shape and size. Volume percentiles were defined as the height at which each percentage point of stem volume was accumulated. Furthermore, absolute and relative volume and tapering below and above 50% of tree height, represented attributes that can be generated from a taper curve. As taper curve has been impractical to measure, these kinds of attributes have not been widely applied and can thus be considered novel in the field of growth and yield. Relative volume below and above 50% of tree height was obtained by dividing the absolute volume of part of a stem with total stem volume. Relative tapering, on the other hand, was produced through division of absolute tapering of bottom and top part of a stem by the diameter at the lowest point, in other words at 0 m and at 50% of tree height for relative tapering of bottom and top part of a stem, respectively. All the attributes were generated for 2174 Scots pine trees within the 27 sample plots.

**Table 3.**
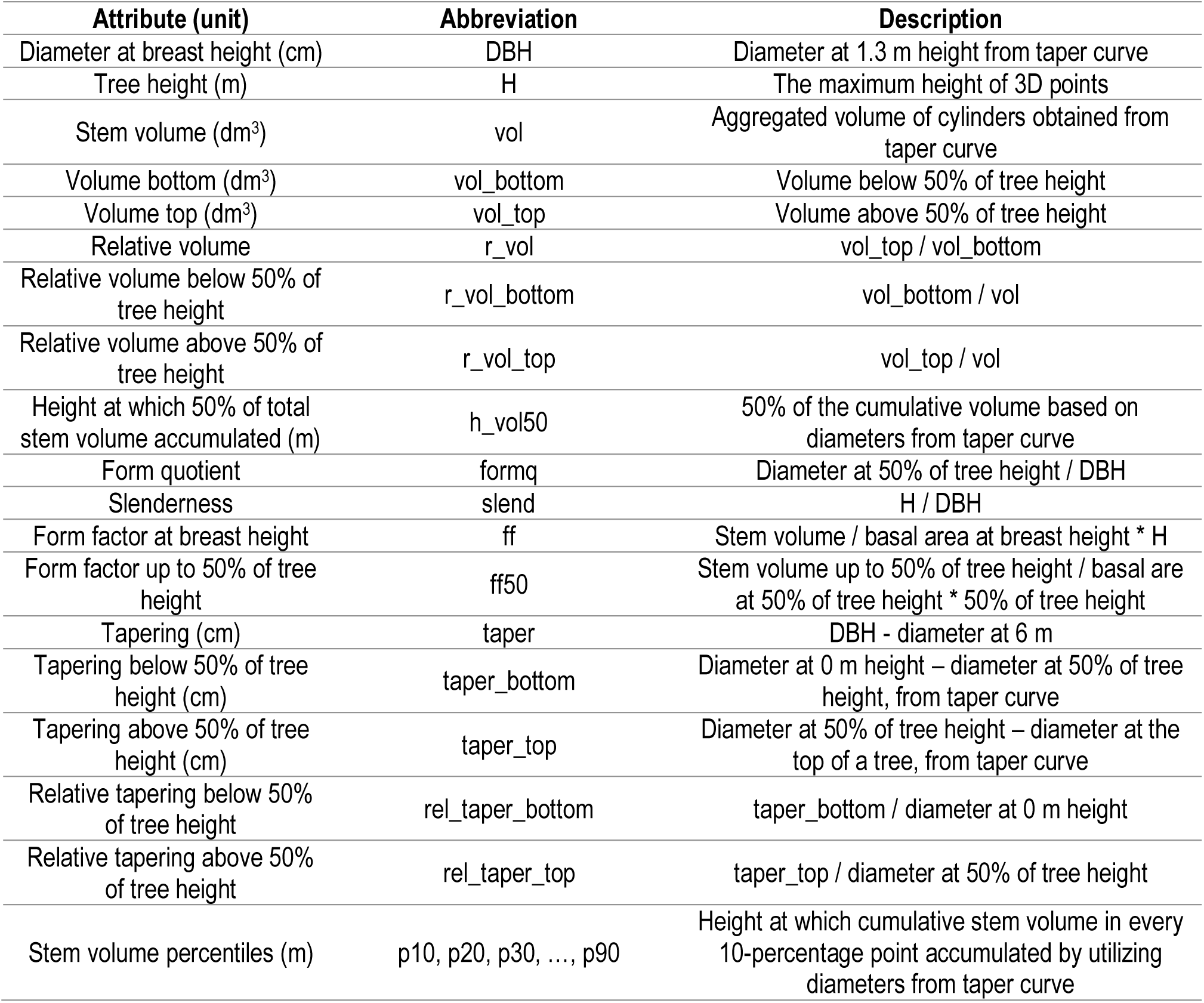
Single tree attributes characterizing stem shape and size derived from taper curves generated with information from terrestrial laser scanning and unmanned aerial vehicles.

| Attribute (unit) | Abbreviation | Description |
| --- | --- | --- |
| Diameter at breast height (cm) | DBH | Diameter at 1.3 m height from taper curve |
| Tree height (m) | H | The maximum height of 3D points |
| Stem volume (dm<sup>3</sup>) | vol | Aggregated volume of cylinders obtained from taper curve |
| Volume bottom (dm<sup>3</sup>) | vol_bottom | Volume below 50% of tree height |
| Volume top (dm<sup>3</sup>) | vol_top | Volume above 50% of tree height |
| Relative volume | r_vol | vol_top / vol_bottom |
| Relative volume below 50% of tree height | r_vol_bottom | vol_bottom / vol |
| Relative volume above 50% of tree height | r_vol_top | vol_top / vol |
| Height at which 50% of total stem volume accumulated (m) | h_vol50 | 50% of the cumulative volume based on diameters from taper curve |
| Form quotient | formq | Diameter at 50% of tree height / DBH |
| Slenderness | slend | H / DBH |
| Form factor at breast height | ff | Stem volume / basal area at breast height * H |
| Form factor up to 50% of tree height | ff50 | Stem volume up to 50% of tree height / basal area at 50% of tree height * 50% of tree height |
| Tapering (cm) | taper | DBH - diameter at 6 m |
| Tapering below 50% of tree height (cm) | taper_bottom | Diameter at 0 m height – diameter at 50% of tree height, from taper curve |
| Tapering above 50% of tree height (cm) | taper_top | Diameter at 50% of tree height – diameter at the top of a tree, from taper curve |
| Relative tapering below 50% of tree height | rel_taper_bottom | taper_bottom / diameter at 0 m height |
| Relative tapering above 50% of tree height | rel_taper_top | taper_top / diameter at 50% of tree height |
| Stem volume percentiles (m) | p10, p20, p30, ..., p90 | Height at which cumulative stem volume in every 10-percentage point accumulated by utilizing diameters from taper curve |

Due to the data structure (i.e. several sample plots in each study site), a nested two-level linear mixed-effects model (Equation 1) was fitted using Restricted Maximum Likelihood included in package nlme (Pinheiro et al. 2016) of the R-software (R Core Team, 2019).

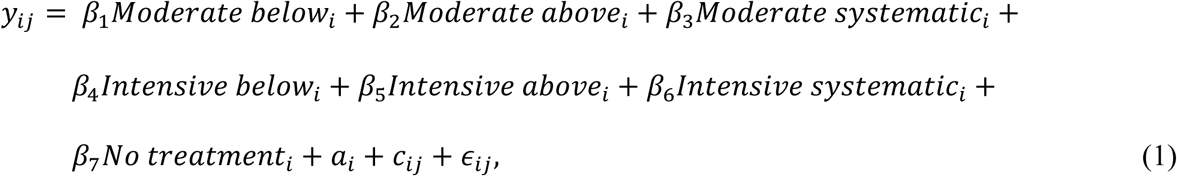

where *y*_*ij*_ is each stem attribute described in Table 3 at a time, *β*_1_, … *β*_7_ are fixed parameters, i, i = 1, …, M, refers to study site, j, j = 1, …,*n*_*i*_, to a plot, *a*_*i*_ and *c*_*ij*_ are normally distributed random effects for sample plot *j* and for sample plot *j* within study site *i*, respectively, with mean zero and unknown, unrestricted variance-covariance matrix, and ϵ_*ij*_ is a residual error with mean zero and unknown variance. The random effects are independent across study sites and sample plots as well as residual errors are independent across trees. The effects of a study site and a sample plot within the study sites on the stem attributes were assessed through their variances.

The analysis of variance was applied in testing the statistically significant difference in the stem attributes affected by the thinning treatments, the study sites as well as the plots within study sites. Furthermore, to reveal the possible statistically significant difference in the stem attributes between a thinning treatment against other treatments, Tukey’s honest significance test was applied.

To compare the taper curves after the different thinning treatments that were carried out in 2005 for study site Palomäki and in 2006 in Pollari and Vesijako, diameters for all trees within each treatment were estimated with a smoothing cubic spline to relative heights of 1%, 2.5%, 5%, 7.5%, 10%, 15%, 20%, …, 95%, and 100% using the taper curve measurements obtained with information from TLS and UAV. This was carried out to normalize the differences in tree height. Finally, a mean taper curve was calculated for each treatment together with its standard deviation.

As especially thinning type implicitly affects the size of the remaining trees (e.g. in thinning from below small trees were removed), we examined the basic statistics (i.e. minimum, mean, maximum, and standard deviation) of relative stem attributes, namely relative volume, relative volume below and above 50% tree height as well as relative tapering below and above 50% of tree height but also slenderness and both form factor at breast height and form factor up to 50% of tree height. These attributes disregard tree size and demonstrate more objectively the effects of different thinning intensity and type and can, therefore, reveal the variation in stem shape and size.

Although taper curve for each tree was only available for one time point, field measurements before and after the thinning treatments were on hand. Post-thinning growth in DBH, height, and stem volume was calculated for all live trees that were in the sample plots during the last measurements. Additionally, difference in slenderness was assessed as a ratio between the first and last measurements to evaluate post-thinning growth. To evaluate if the growth of DBH, height, and stem volume as well as difference in slenderness varied statistically significantly (p-value<0.05) between thinning treatments, a similar nested-two level linear mixed-effects model as presented in Equation 1 was fitted for the change-related attributes. Moreover, similar statistical analyses (i.e. analysis of variance, Tukey’s honest significance test) were utilized in testing difference in these change-related attributes between the thinning treatments. Additionally, difference in ratio between stem volume from the first and last field measurement was also included to disregard the tree size in growth assessments.

## Results

When the effects of a same thinning type (i.e. from below, from above, and systematic) but of different intensity (i.e. moderate and intensive) on the stem attributes characterizing stem shape and size were compared, intensive thinning mainly produced larger attributes characterizing absolute tree size (Figure 3). Similarly, thinning from below (both moderate and intensive) mostly resulted in larger stem attributes characterizing absolute stem size compared to respective attributes caused by thinning from above and systematic with the same intensity (Figure 3). In relative attributes that disregard tree size, such as relative volumes and taperings as well as form factor at breast height, however, the difference between thinning intensity and type was smaller (Figure 3). On the contrary, the difference in slenderness and form factor up to 50% of tree height between thinning intensity and type was visible. There was a noticeable trend between moderate thinning types in stem volume percentiles, in other words in moderate systematic thinning and thinning from above stem volume percentiles were accumulated at lower heights of a stem compared to moderate thinning from below (Figure 4). Between intensive thinning types there was no such trend but intensive thinning from above, among the intensive thinnings, produced the lowest heights for the stem volume percentile accumulation, indicating larger volume accumulation in lower heights of a stem.

**Figure 3.**
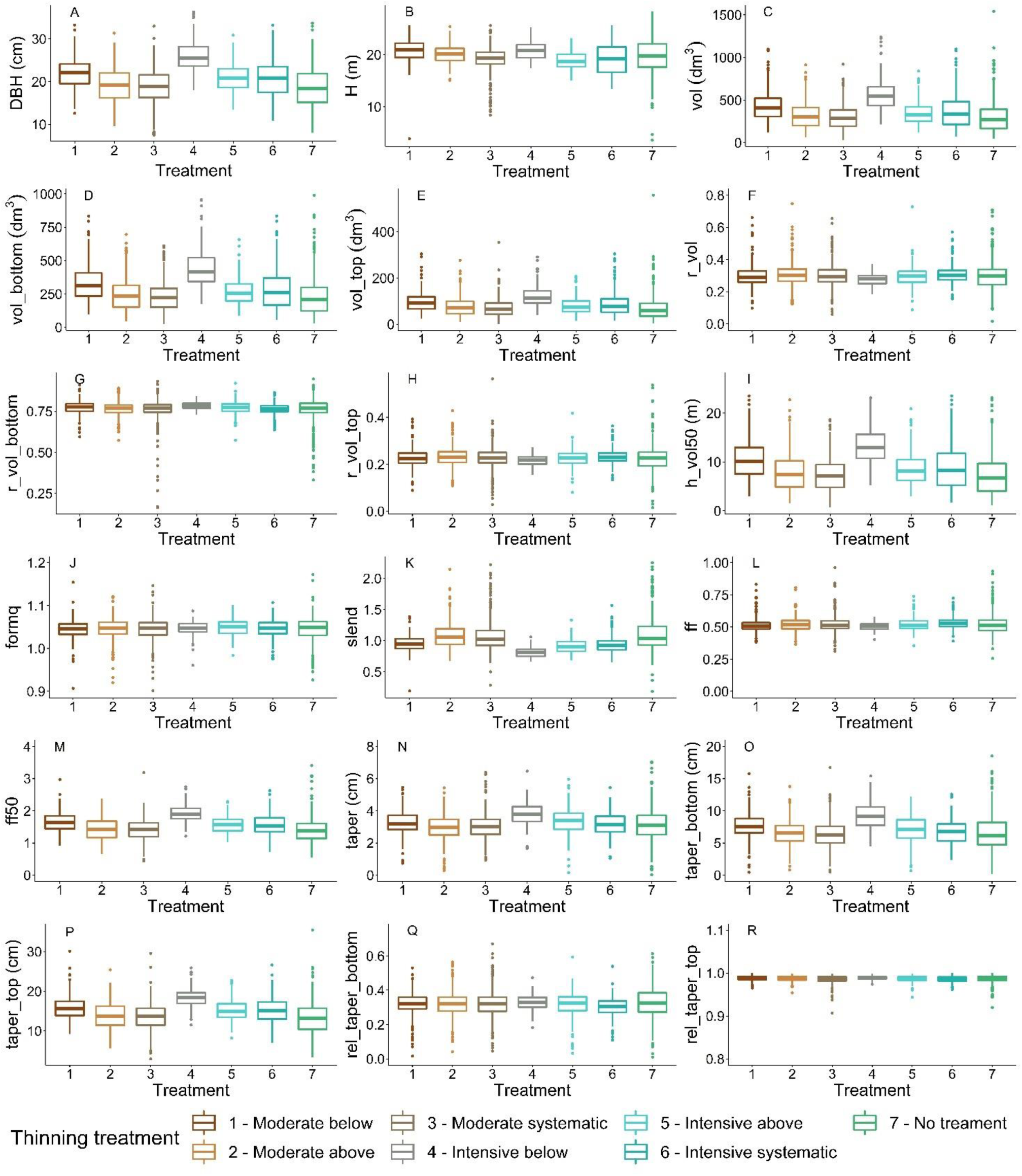
Stem attributes (i.e. diameter at breast height (DBH) (A), tree height (B), stem volume (C), volume below 50% of tree height (vol_bottom) (D), volume above 50% of tree height (vol_top) (E), relative volume (r_vol) (F), relative volume below 50% of tree height (r_vol_bottom) (G), relative volume above 50% of tree height (r_vol_top) (H), height at which 50% of stem volume has accumulated (h_vol50) (I), form quotient (formq) (J), slenderness (slend) (K), form factor at breast height (ff) (L), form factor up to 50% of tree height (ff50) (M), tapering (taper) (N), tapering below 50% of tree height (taper_bottom) (O), tapering above 50% of tree height (taper_top) (P), relative tapering below 50% of tree height (rel_taper_bottom) (Q), and relative tapering above 50% of tree height (rel_taper_top) (R)) caused by seven thinning treatments implemented in 2005-2006. Please refer to Table 3 for the descriptions of the attributes.

**Figure 4.**
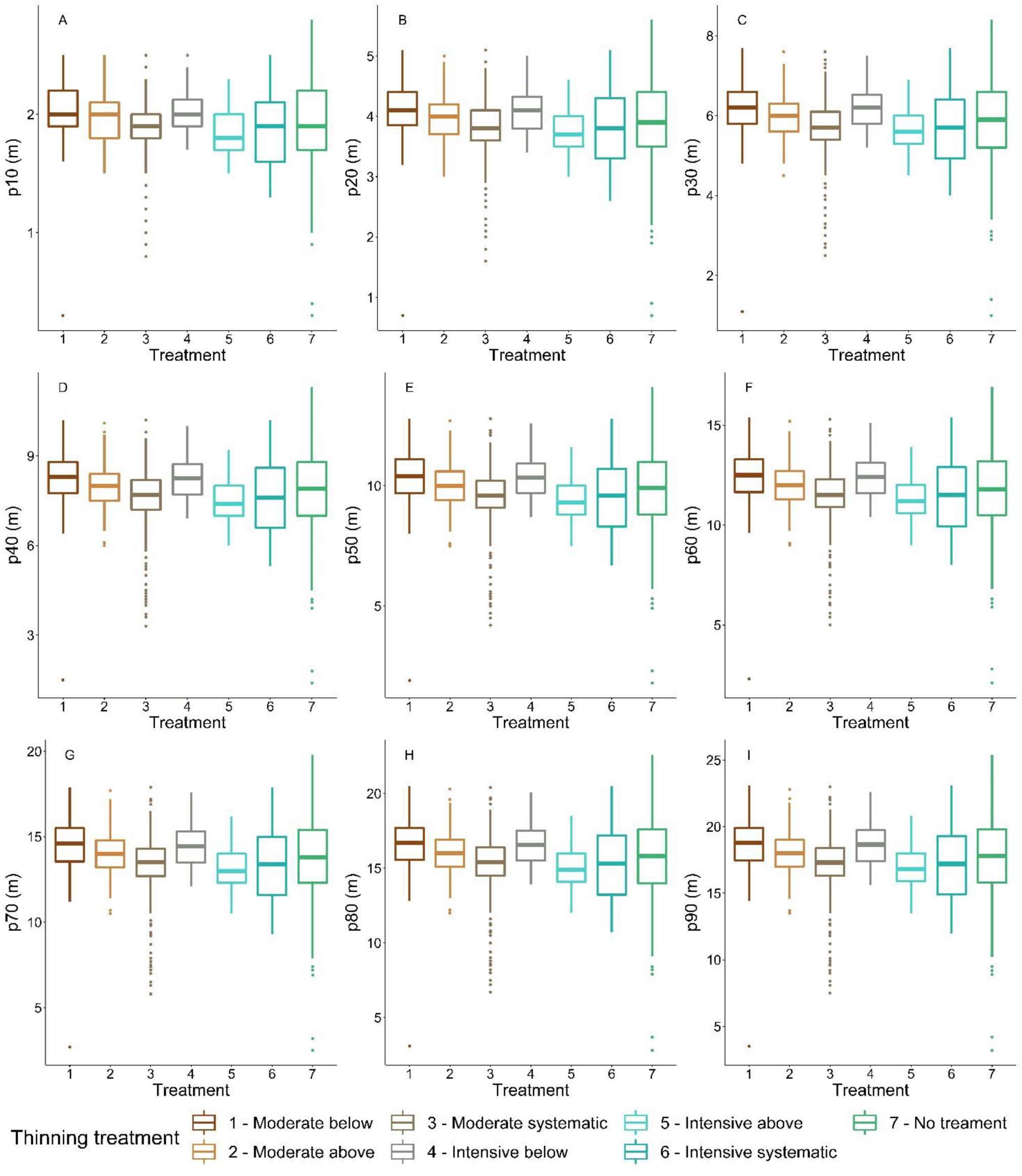
Differences caused by thinning treatments in 2005-2006 and growth of stem volume percentiles (i.e. height at which percentiles of volume accumulated; 10^th^: A, 20^th^: B, 30^th^: C, 40^th^: D, 50^th^: E, 60^th^: F, 70^th^: G., 80^th^: H, and 90^th^: I).

The nested two-level linear mixed-effects models provided quantitative details about differences in stem attributes between thinning treatments depicted in Figures 3 and 4 but also whether their effects were statistically significant (Table S1). Variances of the random part of the mixed-effects model explained the variation in the stem attributes between the study sites and the sample plots within a study site. They were rather similar for all other stem attributes except for form quotient for which the variance of a study site was a ten of times larger (3.128^2^) than the plot-level variance (0.004^2^) (Table S2). Furthermore, for relative volume (i.e. ratio between volume below and above 50% of tree height) the plot-level variance was considerably larger (6.326^2^) than the variance at study site level (0.014^2^) similar to tapering above 50% of tree height (plot level 1.464^2^ vs study site 0.001^2^). These suggest that most of the variation between the stem attributes was explained by both a study site and a plot within a study site, but a study site affected considerably more on form quotient whereas a sample plot affected considerably more on relative volume and (absolute) tapering above 50% of tree height.

As the differences were not noticeable for all stem attributes between thinning treatments in Figures 3 and 4, the analysis of variance only revealed statistically significant differences (p-value ≤ 0.05) in DBH, volume, volume below and above 50% of tree height, relative volume, relative volume below 50% of tree height, height at which 50% of total stem volume accumulated, slenderness, form factor up to 50% of tree height, tapering as well as tapering below and above 50% of tree height. However, it did not reveal between which treatments the differences occurred. Tukey’s honest significance test, however, showed statistically significant difference (p-value ≤ 0.05) between thinning treatments (Table S3). Furthermore, the analysis of variance showed that there was a significant difference (p-value ≤ 0.05) between the three study sites in all stem attributes except form quotient and relative tapering above 50% of tree height. When the analysis of variance was used for assessing difference between plots within a study site, there was statistically significant difference (p-value ≤ 0.05) in all stem attributes except relative volume and relative volume below 50% of tree height.

When comparing the influence of thinning intensity on the relative stem attributes, only relative volume and relative volume above 50% of tree height differed significantly (p-value ≤ 0.05) between moderate thinning from above and intensive thinning from below. The thinning type within the intensive thinnings affected relative volume and relative volume above 50% of tree height from the relative stem attributes; they differed between intensive from below and systematic. However, statistically significant difference was consistently found in slenderness when comparing both thinning intensity and type (Table S3) whereas form factor up to 50% of tree height differed statistically significantly between thinning types of intensive thinning.

A taper curve depicts stem shape and size and a mean taper curve for each treatment with its standard deviation is presented in Figure 5. Larger diameters are visible for taper curves of intensive thinnings compared to moderate thinnings, in other words more volume and through that biomass can be expected to be allocated in stems in intensively thinned plots. Similarly, when the thinning from below and above are compared, larger diameters are observed in the bottom part of a stem (i.e., below 50% relative height) in thinning from below indicating larger absolute bottom parts. Furthermore, diameters in the top part of the mean taper curves (i.e., above 50% of relative height) are more similar between thinning treatments. Tapering in the bottom and top parts of a stem was smaller in thinnings from above compared to thinnings from below (with both intensities) (Figure 3).

**Figure 5.**
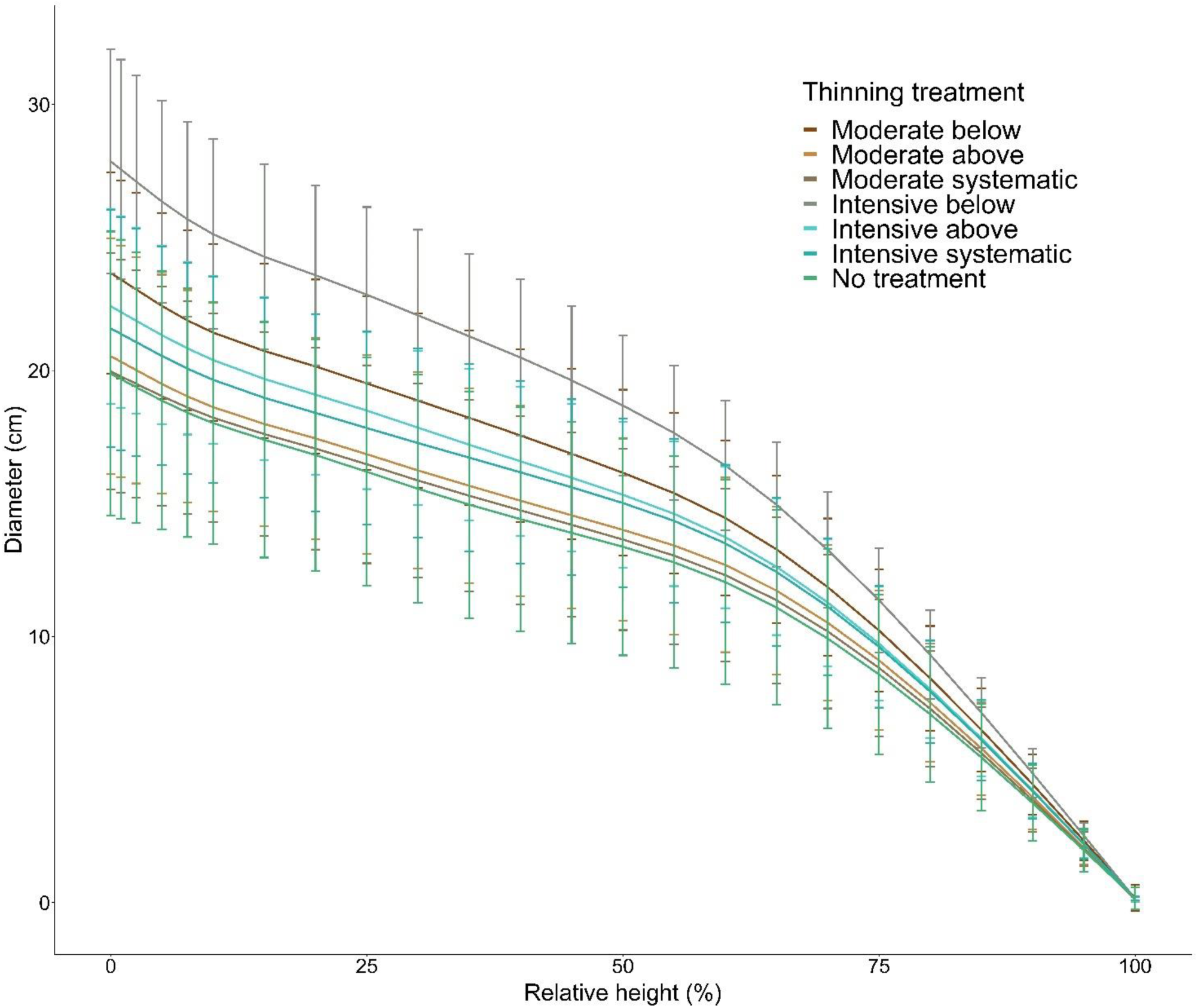
Mean taper curves with standard deviation (error bars) of each thinning treatment for relative heights.

The relative volume and tapering attributes as well as slenderness and form factors were assessed separately through basic statistics to examine the variation due to the thinning treatments. The relative attributes do not consider tree size that is implicitly included in thinning type (i.e. tree size affects which trees are removed). There were no clear differences between thinning treatments in relative volumes (i.e. relative volume, relative volume below and above 50% of tree height) or taperings (Figure 3) and statistically significant difference was only found in relative volume and relative volume above 50% of tree height (Tables 4 and S3). The intensive thinnings (i.e. below, above, and systematic) mainly resulted in smaller standard deviation compared to the relative volume and tapering attributes of their corresponding moderate counterparts (Table 4). Only between moderate and intensive thinning from above the standard deviation of relative tapering above 50% of tree height was very similar, namely 0.006 and 0.007, respectively. When the thinning type was compared, thinning from below produced smaller standard deviation than thinning from above and systematic thinning. Intensive thinning from below produced the smallest standard deviation in all the attributes characterizing relative volume and tapering whereas it was the largest in control plots (i.e. No treatment). However, when especially considering slenderness there was statistically significant difference when comparing different thinning intensity of the same thinning type as well as when thinning type within the same thinning intensity was compared (Tables 4 and S3). As slenderness is also defined as a ratio between tree height and DBH, the difference in it indicates varying growth response between DBH and height after different thinning treatments. Intensive thinning (i.e. below, above, and systematic) as well as moderate below resulted in slenderness below 1 indicating larger DBH compared to tree height which suggest more allocation of post-thinning growth in diameters than in height. Form factor up to 50% of tree height, on the other hand, characterizes shape of the bottom part of a stem (i.e. below 50% of tree height) and it was statistically significantly different between thinning types of intensive thinning (Tables 4 and S3). This suggests that thinning type could also affect the shape of the bottom part of a stem. Intensive thinning and thinning from below also produced the smallest variation (i.e. standard deviation) for slenderness whereas the standard deviation of both form factor at breast height and form factor up to 50% of tree height was not as consistent between thinning types (Table 4).

**Table 4.**
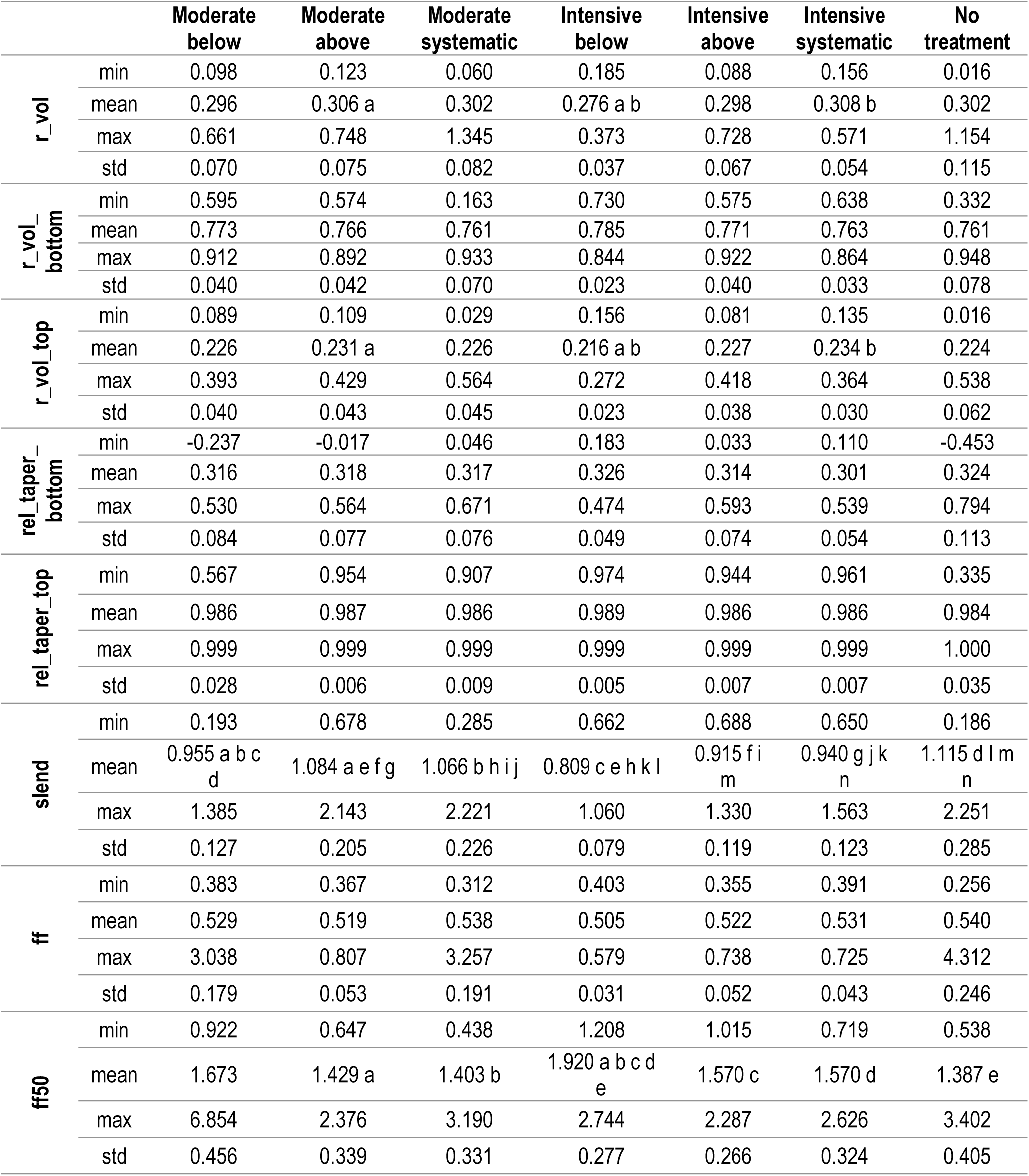
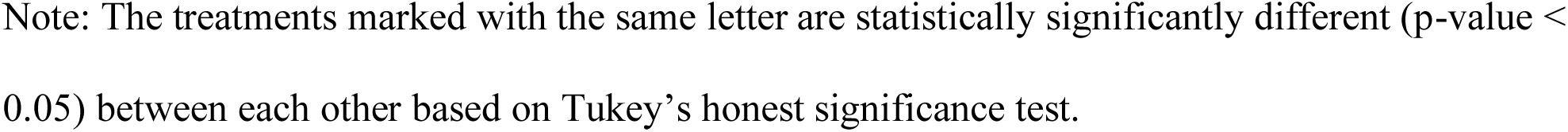
Minimum, mean, maximum and standard deviation (std) of relative stem attributes namely relative volume (r_vol), relative volume below 50% of tree height (r_vol_bottom), relative volume above 50% of tree height (r_vol_top), relative tapering below 50% of tree height (rel_taper_bottom), relative tapering above 50% of tree height (rel_taper_top), slenderness (slend), form factor at breast height (ff), and form factor up to 50% of tree height (ff50) between thinning treatments. Please refer to Table 3 for the descriptions of the attributes.

|  |  | Moderate<br>below | Moderate<br>above | Moderate<br>systematic | Intensive<br>below | Intensive<br>above | Intensive<br>systematic | No<br>treatment |
| --- | --- | --- | --- | --- | --- | --- | --- | --- |
| r_vol | min | 0.098 | 0.123 | 0.060 | 0.185 | 0.088 | 0.156 | 0.016 |
|  | mean | 0.296 | 0.306 a | 0.302 | 0.276 a b | 0.298 | 0.308 b | 0.302 |
|  | max | 0.661 | 0.748 | 1.345 | 0.373 | 0.728 | 0.571 | 1.154 |
|  | std | 0.070 | 0.075 | 0.082 | 0.037 | 0.067 | 0.054 | 0.115 |
| r_vol_bottom | min | 0.595 | 0.574 | 0.163 | 0.730 | 0.575 | 0.638 | 0.332 |
|  | mean | 0.773 | 0.766 | 0.761 | 0.785 | 0.771 | 0.763 | 0.761 |
|  | max | 0.912 | 0.892 | 0.933 | 0.844 | 0.922 | 0.864 | 0.948 |
|  | std | 0.040 | 0.042 | 0.070 | 0.023 | 0.040 | 0.033 | 0.078 |
| r_vol_top | min | 0.089 | 0.109 | 0.029 | 0.156 | 0.081 | 0.135 | 0.016 |
|  | mean | 0.226 | 0.231 a | 0.226 | 0.216 a b | 0.227 | 0.234 b | 0.224 |
|  | max | 0.393 | 0.429 | 0.564 | 0.272 | 0.418 | 0.364 | 0.538 |
|  | std | 0.040 | 0.043 | 0.045 | 0.023 | 0.038 | 0.030 | 0.062 |
| rel_taper_bottom | min | -0.237 | -0.017 | 0.046 | 0.183 | 0.033 | 0.110 | -0.453 |
|  | mean | 0.316 | 0.318 | 0.317 | 0.326 | 0.314 | 0.301 | 0.324 |
|  | max | 0.530 | 0.564 | 0.671 | 0.474 | 0.593 | 0.539 | 0.794 |
|  | std | 0.084 | 0.077 | 0.076 | 0.049 | 0.074 | 0.054 | 0.113 |
| rel_taper_top | min | 0.567 | 0.954 | 0.907 | 0.974 | 0.944 | 0.961 | 0.335 |
|  | mean | 0.986 | 0.987 | 0.986 | 0.989 | 0.986 | 0.986 | 0.984 |
|  | max | 0.999 | 0.999 | 0.999 | 0.999 | 0.999 | 0.999 | 1.000 |
|  | std | 0.028 | 0.006 | 0.009 | 0.005 | 0.007 | 0.007 | 0.035 |
| slend | min | 0.193 | 0.678 | 0.285 | 0.662 | 0.688 | 0.650 | 0.186 |
|  | mean | 0.955 a b c d | 1.084 a e f g | 1.066 b h i j | 0.809 c e h k l | 0.915 f i m | 0.940 g j k n | 1.115 d l m n |
|  | max | 1.385 | 2.143 | 2.221 | 1.060 | 1.330 | 1.563 | 2.251 |
|  | std | 0.127 | 0.205 | 0.226 | 0.079 | 0.119 | 0.123 | 0.285 |
| ff | min | 0.383 | 0.367 | 0.312 | 0.403 | 0.355 | 0.391 | 0.256 |
|  | mean | 0.529 | 0.519 | 0.538 | 0.505 | 0.522 | 0.531 | 0.540 |
|  | max | 3.038 | 0.807 | 3.257 | 0.579 | 0.738 | 0.725 | 4.312 |
|  | std | 0.179 | 0.053 | 0.191 | 0.031 | 0.052 | 0.043 | 0.246 |
| ff50 | min | 0.922 | 0.647 | 0.438 | 1.208 | 1.015 | 0.719 | 0.538 |
|  | mean | 1.673 | 1.429 a | 1.403 b | 1.920 a b c d e | 1.570 c | 1.570 d | 1.387 e |
|  | max | 6.854 | 2.376 | 3.190 | 2.744 | 2.287 | 2.626 | 3.402 |
|  | std | 0.456 | 0.339 | 0.331 | 0.277 | 0.266 | 0.324 | 0.405 |

Note: The treatments marked with the same letter are statistically significantly different (p-value < 0.05) between each other based on Tukey's honest significance test.

Growth in DBH, height, and stem volume was calculated based on the first and last field measurements (Table 5). Additionally, difference in slenderness between the first and last field measurements was assessed. The largest growth in DBH and stem volume was found in the intensive thinning from below, 7.0 cm and 314.6 dm^3^, respectively, whereas the largest height growth (i.e. 5.2 was observed in plots without thinning treatment. For the moderate thinnings, difference in slenderness ratio was <1 whereas for the intensive thinnings it was >1 indicating smaller difference in height-diameter ratio in denser plots. The analysis of variance showed statistically significant difference (p-value ≤ 0.05) in growth of DBH and volume as well as in difference in slenderness between thinning treatments. Statistically significant difference (p-value ≤ 0.05) between study sites, on the other hand, was found in growth of DBH and height as well as in difference in slenderness whereas difference in all these change-related attributes was statistically significant between plots within study sites.. In DBH growth and difference in slenderness significant difference (p-value ≤ 0.05) was found between thinning intensity whereas significant difference (p-value ≤ 0.05) in volume growth was found between thinning type (intensive thinning treatments). When ratio of stem volume between the first and last field measurements was assessed, the significant difference (p-value ≤ 0.05) was found between thinning intensity and type but only between intensive systematic thinning.and intensive thinnings from below and above.

**Table 5.**
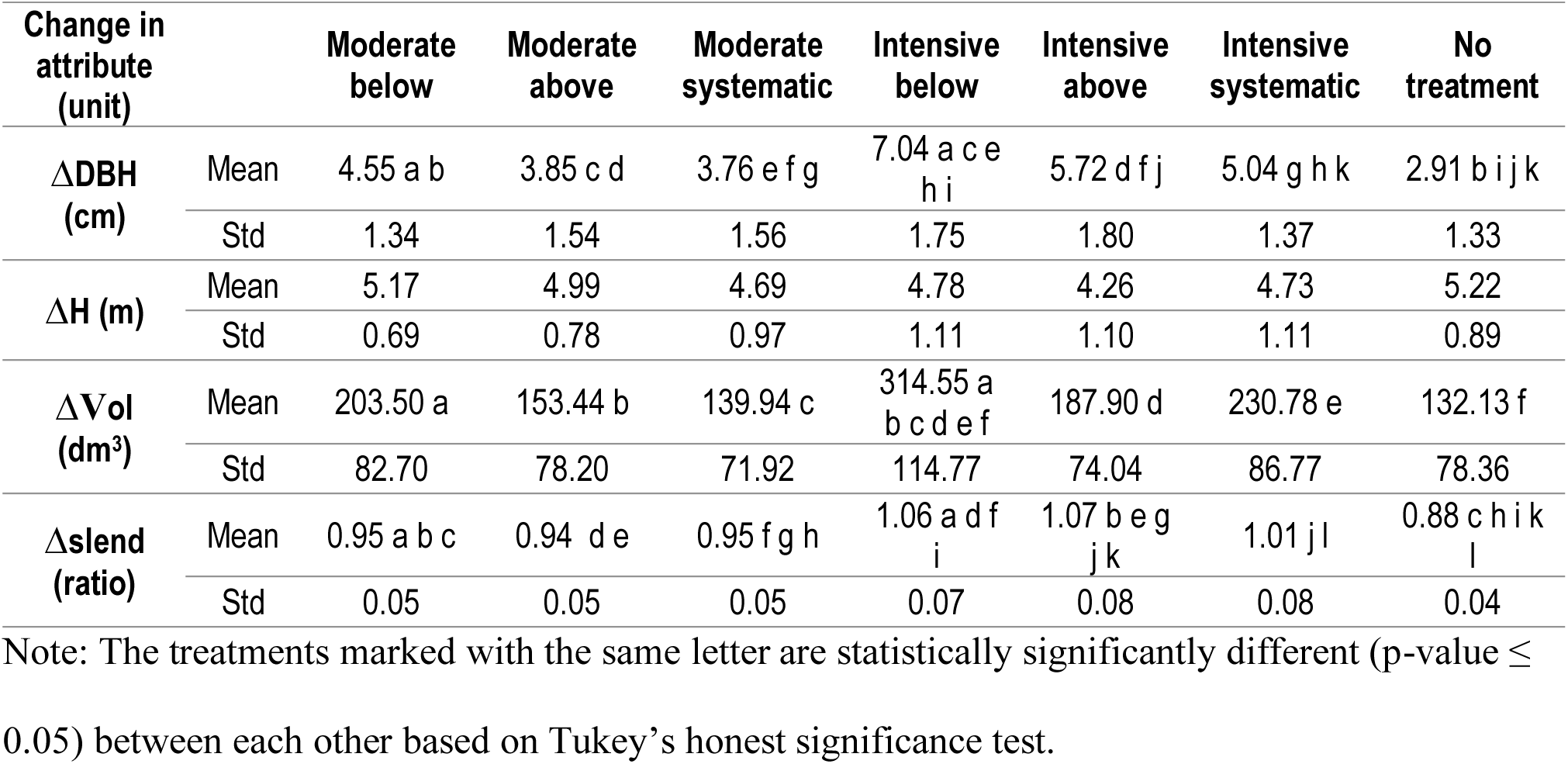
Mean growth with standard deviation (Std) in diameter at breast height (ΔDBH), height (ΔH), and stem volume (ΔVol) as well as mean difference with standard deviation in slenderness ratio (Δslend) between first (2005-2006) and last (2018-2019) field measurements.

## Discussion

The objective of this work was to examine how thinning intensity but also thinning type (i.e. from below, from above, and systematic) influenced stem shape and size of Scots pine trees. The results showed that there was no clear difference in relative stem attributes characterizing stem shape and size although in absolute stem attributes difference was found between thinning type (research question 1). However, the relationship between DBH and height (i.e. slenderness) differed between both thinning intensity and type (research question 2). Finally, when assessing the similarity of the relative attributes between different thinning treatments, intensive thinning, and especially intensive thinning from below led to the smallest variation in those attributes (research question 3). Thus, our hypothesis about thinning intensity and type affecting shape and size of Scots pine trees can be accepted, although no clear difference in most of the relative stem attributes was observed. Difference in slenderness (i.e. growth allocation between DBH and tree height), however, affects both stem shape and size.

Taper curves derived from the 3D information provided by the TLS and UAV enabled generating new attributes characterizing stem shape and size that have formerly only been available through destructive measurements and thus provided novel and quantitative information about stem shape and size (in both absolute and relative scale) for forest management and ecology. As stem size is inherently included in the thinning treatments, there is already variation in the absolute size of the remaining trees. With intensive thinning, more trees were removed that ensured more resources for the remaining trees whereas with moderate thinning the resultant tree density was larger. Smaller trees were removed in thinnings from below and larger trees were left to grow whereas the situation was the opposite in thinnings from above and systematically. Thus, in addition to different growth effects, our findings about differences in absolute stem size between these thinning treatments is expected. Conversely, there was no apparent distinction in most of the realization of the (absolute) stem attributes between thinning from above and systematic with either moderate or intensive intensity. This is most probably due to the similarity between thinning from above and systematic thinning that both concentrated on removing larger trees. The relative attributes, on the other hand, could enhance the assessment of the effects of thinning treatments as absolute tree size was disregarded. Many earlier studies have concentrated on differences in absolute size affected by different thinning treatments (Juodvalkis et al. 2005, Eriksson 2006, Nilsson et al. 2010, Valinger et al. 2019). Nilsson et al. (2010) reported no difference in stand-level stem volume between thinning from below and above for Scots pine. We only considered tree-level stem volume but found a significant difference in it between thinning type (intensive below vs intensive above, intensive below and intensive systematic, and intensive below vs no treatment). Valinger et al. (2019) found no effect of thinning on tree height growth which is similar to our findings. Mäkinen et al. (2005) compared absolute tapering between different thinning intensities and found larger tapering when thinning intensity increased which is in line with our results. Montero et al. (2001) as well as Mäkinen & Isomäki (2004c) and Mäkinen et al. (2005), on the other hand, considered difference in slenderness when thinning intensity varied in Scots pine stands. They all reported smaller slenderness when thinning intensity increased in Scots pine stands and it was corroborated by this study.

Del Rio et al. (2017) reviewed thinning experiments of Scots pine and pointed out that in addition to effects on growth and yield, thinning also affects stand stability against snow and wind and for that attributes with the greatest importance are tree height, height and DBH ratio (i.e. slenderness), and the length of a living crown. Light thinning from below in early stage of forest development improved the resistance to wind similarly to heavy thinning from below in early stage also improved the resistance to snow. Moreover, heavy thinning increased tree growth recovery as a response to drought. From biodiversity perspective, del Rio et al. (2017) summarized that heavy thinning enhanced understory vegetation but decreased structural diversity whereas thinning in general had positive effects on the provision of ecosystem services such as wood quality and could also positively affect the yield of berries and mushrooms.

Stem attributes have been generated from TLS-based point clouds to study the effects of thinning in Central Europe. Jacobs et al. (2019) reported smaller form factor with less competition whereas tapering in butt logs decreased when competition increased for Norway spruce in southern Germany. Our results did not reveal notable difference in form factor between moderate and intensive thinning whereas tapering below 50% of tree height was smaller for moderate thinnings than for intensive thinnings corroborating results by Jacobs et al. (2019). Georgi et al. (2018), on the other hand, found significant difference in tapering generated from TLS data between unmanaged and intensively managed beech stands. Most of their attributes were related to crown dimensions or height of branches similar to the study by Juchheim et al. (2017b) who concentrated on crown expansion in both horizontal and vertical directions. In addition to stem shape and size, long-term growth and yield experiments have shown that thinning treatments affect crown ratio (Mäkinen et al. 2005), crown area increment (Juodvalkis et al. 2005), as well as biofuel production and standing biomass (Eriksson 2006). TLS has been used in providing tree biomass of conifers (Hauglin et al. 2012, Kankare et al. 2013) and deciduous (Srinivasan et al. 2014, Stovall et al. 2018) trees as well as in the tropics (Calders et al. 2015, Gonzalez de Tanago et al. 2017). It also provides a means for measuring crown dimensions and structure (Fernández-Sarría et al. 2013, Metz et al. 2015, Ferrarese et al. 2015) that can be further used in assessing structural complexity (Seidel 2017) but also effects of management (Juchheim 2017a, Stiers et al. 2018) and species composition (Bayer et al. 2013, Seidel et al. 2016, Barbeito et al. 2017) on tree architecture. However, there is still a limited understanding of the holistic effects of thinning intensity and type on tree architecture (e.g. crown dimension and structure) as well as structural complexity in boreal forest conditions in which TLS can be of an assist. Furthermore, it was assumed that 3D information also from above the canopy (i.e. from the UAV) would enhance information on tree height and therefore the reliability of the taper curves. This assumption can be accepted as Yrttimaa et al. (2020) showed that the combination of UAV and TLS point clouds can improve reliability of estimates for mean height at both stand and tree level.

Intensive thinning expanded the growing space of and reduced the competition between the remaining trees allowing them to allocate the photosynthesis products not just sustaining respiration and existing parts but also for wood formation and growth. Correspondingly, thinning from below enhanced the growing conditions of dominant and co-dominant trees and enabled trees that were already in advanced position to boost their development. The relationship between DBH and tree height (i.e. slenderness) varied between both thinning intensity and type indicating different strategy for the allocation of photosynthesis products between Scots pine trees after different thinning treatments.

## Conclusions

Taper curve characterizes stem dimensions, tapering, shape, and especially volume allocation of an individual tree. Therefore, point clouds from TLS and UAV were utilized in reconstructing stems and deriving taper curves for Scots pine trees growing in similar site conditions (e.g., same fertility based on vegetation) but with varying management history enabling us to investigate the effects of forest management (i.e. thinning) on stem shape and size. Our results showed that more intensive thinning than usually applied in Finland caused larger Scots pine trees in size (i.e., DBH and volume) compared to moderate thinning (i.e. the state-of-the art thinning intensity). When stem shape and size were compared between thinning type (i.e., from below, from above, or systematic), thinning from below resulted in larger bottom part of a stem (i.e. absolute volume). A large bottom part and large Scot pine trees in total could enhance wood production and especially recovery of saw logs. Quality aspects (e.g. number and quality of branches) were, however, not considered here and they might affect possible economic advantages obtainable from the increased dimensions of Scots pine trees as a result of intensive thinning. Intensive thinning could also enhance adaptation to drought as well as resilience to snow damage of Scots pine trees in Finland.

It can also be concluded that different thinning treatments caused differing relationship between DBH and tree height indicating distinct growth response of Scots pine trees when intensive thinning or thinning from below is applied. Those thinning treatments led to more growing space for the remaining trees and they could allocate their growth in stem instead of accessing light through height growth. There was no clear difference in the stem attributes characterizing relative volume and tapering. They have not been, however, widely used in earlier studies and would thus warrant more investigations from Scots pine trees from around Finland and internationally. Additionally, exploring how other tree species (e.g. Norway spruce) respond after various thinning treatments when assessed with those attributes would further be required. The point clouds from TLS and UAV automatically provided detailed information for stem shape and size from more than 2100 Scots pines enabling creation of novel attributes characterizing post-thinning stem development. The hypothesis that thinning intensity and type affect shape and size of Scots pine trees was accepted as it was shown that intensive thinning and thinning from below produced larger bottom part of stems as well as more similar (i.e. smaller variance) stem shape and size compared to moderate thinning or thinning from above.

## Acknowledgements

The study was funded through the post-doctoral project “The effects of stand dynamics on tree architecture of Scots pine trees” (project number 315079), the Centre of Excellence in Laser Scanning Research (project number 272195), and the project “Autonomous tree health analyzer based on imaging UAV spectrometry” (project number 327861) by the Academy of Finland.

## Supplementary material

**Table S1.**
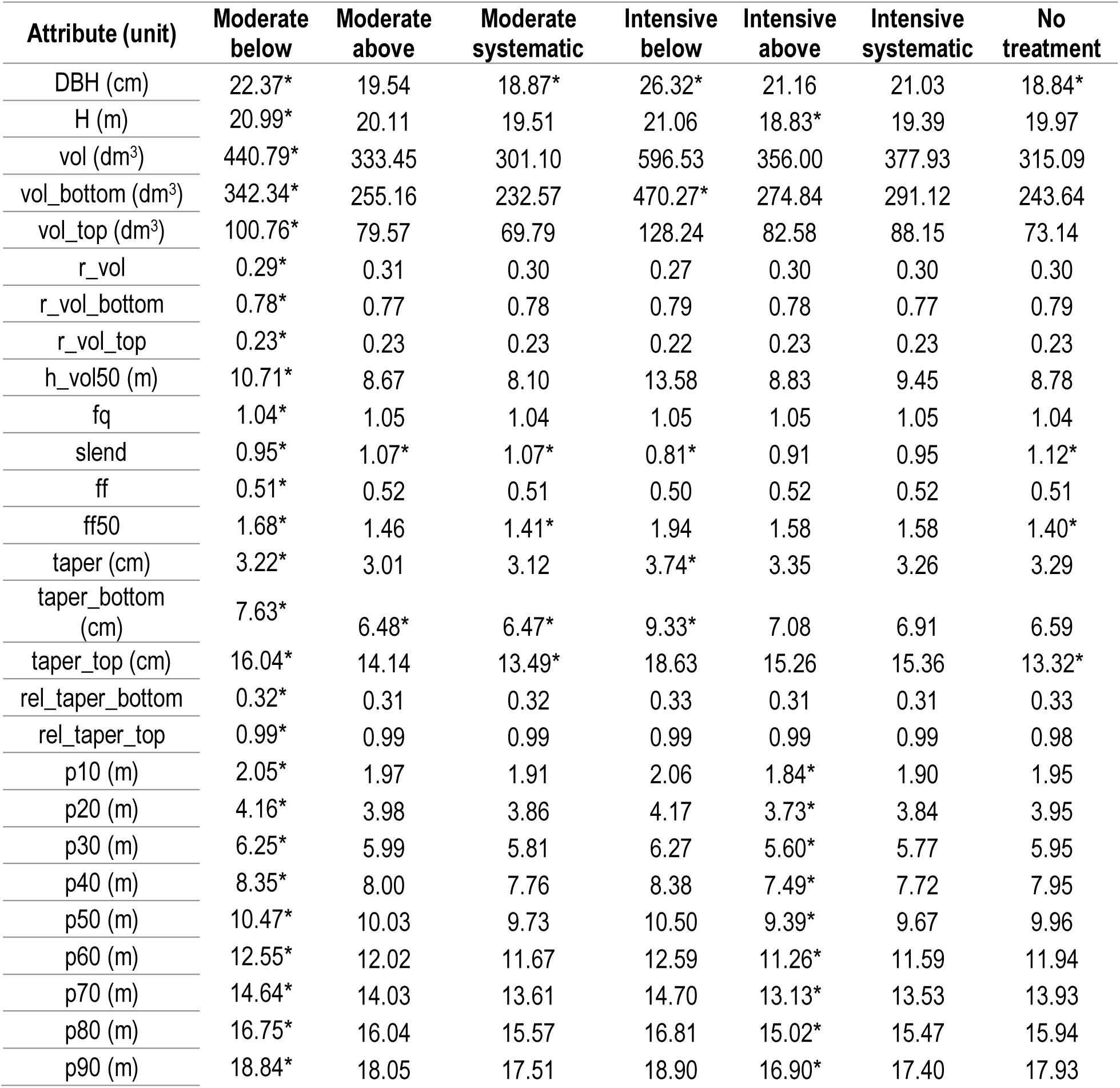
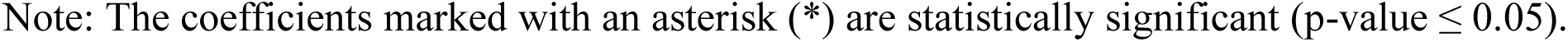
Coefficients of the fixed part of the nested two-level linear mixed-effects model (Equation 1) for each stem attribute. DBH = diameter at breast height, H= tree height, vol = total stem volume, vol_bottom = volume below 50% of tree height, vol_top = volume above 50% of tree height, r_vol = relative volume, r_vol_bottom = relative volume below 50% of tree height, r_vol_top = relative volume above 50% of tree height, h_vol50 = height at which 50% of stem volume accumulated, fq = form quotient, slend = slenderness, ff = form factor at breast height, ff50 = form factor up to 50% of tree height, taper = tapering, taper_bottom = tapering below 50% stem height, taper_top = tapering above 50% of tree height, rel_taper_bottom = relative taper below 50% of tree height, rel_taper_top = relative taper above 50% of tree height, p10,…p90 = height percentiles at which 10%, …90% of stem volume accumulated. Please refer to Table 3 for descriptions of the attributes.

| Attribute (unit) | Moderate<br>below | Moderate<br>above | Moderate<br>systematic | Intensive<br>below | Intensive<br>above | Intensive<br>systematic | No<br>treatment |
| --- | --- | --- | --- | --- | --- | --- | --- |
| DBH (cm) | 22.37* | 19.54 | 18.87* | 26.32* | 21.16 | 21.03 | 18.84* |
| H (m) | 20.99* | 20.11 | 19.51 | 21.06 | 18.83* | 19.39 | 19.97 |
| vol (dm<sup>3</sup>) | 440.79* | 333.45 | 301.10 | 596.53 | 356.00 | 377.93 | 315.09 |
| vol_bottom (dm<sup>3</sup>) | 342.34* | 255.16 | 232.57 | 470.27* | 274.84 | 291.12 | 243.64 |
| vol_top (dm<sup>3</sup>) | 100.76* | 79.57 | 69.79 | 128.24 | 82.58 | 88.15 | 73.14 |
| r_vol | 0.29* | 0.31 | 0.30 | 0.27 | 0.30 | 0.30 | 0.30 |
| r_vol_bottom | 0.78* | 0.77 | 0.78 | 0.79 | 0.78 | 0.77 | 0.79 |
| r_vol_top | 0.23* | 0.23 | 0.23 | 0.22 | 0.23 | 0.23 | 0.23 |
| h_vol50 (m) | 10.71* | 8.67 | 8.10 | 13.58 | 8.83 | 9.45 | 8.78 |
| fq | 1.04* | 1.05 | 1.04 | 1.05 | 1.05 | 1.05 | 1.04 |
| slend | 0.95* | 1.07* | 1.07* | 0.81* | 0.91 | 0.95 | 1.12* |
| ff | 0.51* | 0.52 | 0.51 | 0.50 | 0.52 | 0.52 | 0.51 |
| ff50 | 1.68* | 1.46 | 1.41* | 1.94 | 1.58 | 1.58 | 1.40* |
| taper (cm) | 3.22* | 3.01 | 3.12 | 3.74* | 3.35 | 3.26 | 3.29 |
| taper_bottom<br>(cm) | 7.63* | 6.48* | 6.47* | 9.33* | 7.08 | 6.91 | 6.59 |
| taper_top (cm) | 16.04* | 14.14 | 13.49* | 18.63 | 15.26 | 15.36 | 13.32* |
| rel_taper_bottom | 0.32* | 0.31 | 0.32 | 0.33 | 0.31 | 0.31 | 0.33 |
| rel_taper_top | 0.99* | 0.99 | 0.99 | 0.99 | 0.99 | 0.99 | 0.98 |
| p10 (m) | 2.05* | 1.97 | 1.91 | 2.06 | 1.84* | 1.90 | 1.95 |
| p20 (m) | 4.16* | 3.98 | 3.86 | 4.17 | 3.73* | 3.84 | 3.95 |
| p30 (m) | 6.25* | 5.99 | 5.81 | 6.27 | 5.60* | 5.77 | 5.95 |
| p40 (m) | 8.35* | 8.00 | 7.76 | 8.38 | 7.49* | 7.72 | 7.95 |
| p50 (m) | 10.47* | 10.03 | 9.73 | 10.50 | 9.39* | 9.67 | 9.96 |
| p60 (m) | 12.55* | 12.02 | 11.67 | 12.59 | 11.26* | 11.59 | 11.94 |
| p70 (m) | 14.64* | 14.03 | 13.61 | 14.70 | 13.13* | 13.53 | 13.93 |
| p80 (m) | 16.75* | 16.04 | 15.57 | 16.81 | 15.02* | 15.47 | 15.94 |
| p90 (m) | 18.84* | 18.05 | 17.51 | 18.90 | 16.90* | 17.40 | 17.93 |

Note: The coefficients marked with an asterisk (\*) are statistically significant ( $p\text{-value} \leq 0.05$ ).

**Table S2.**
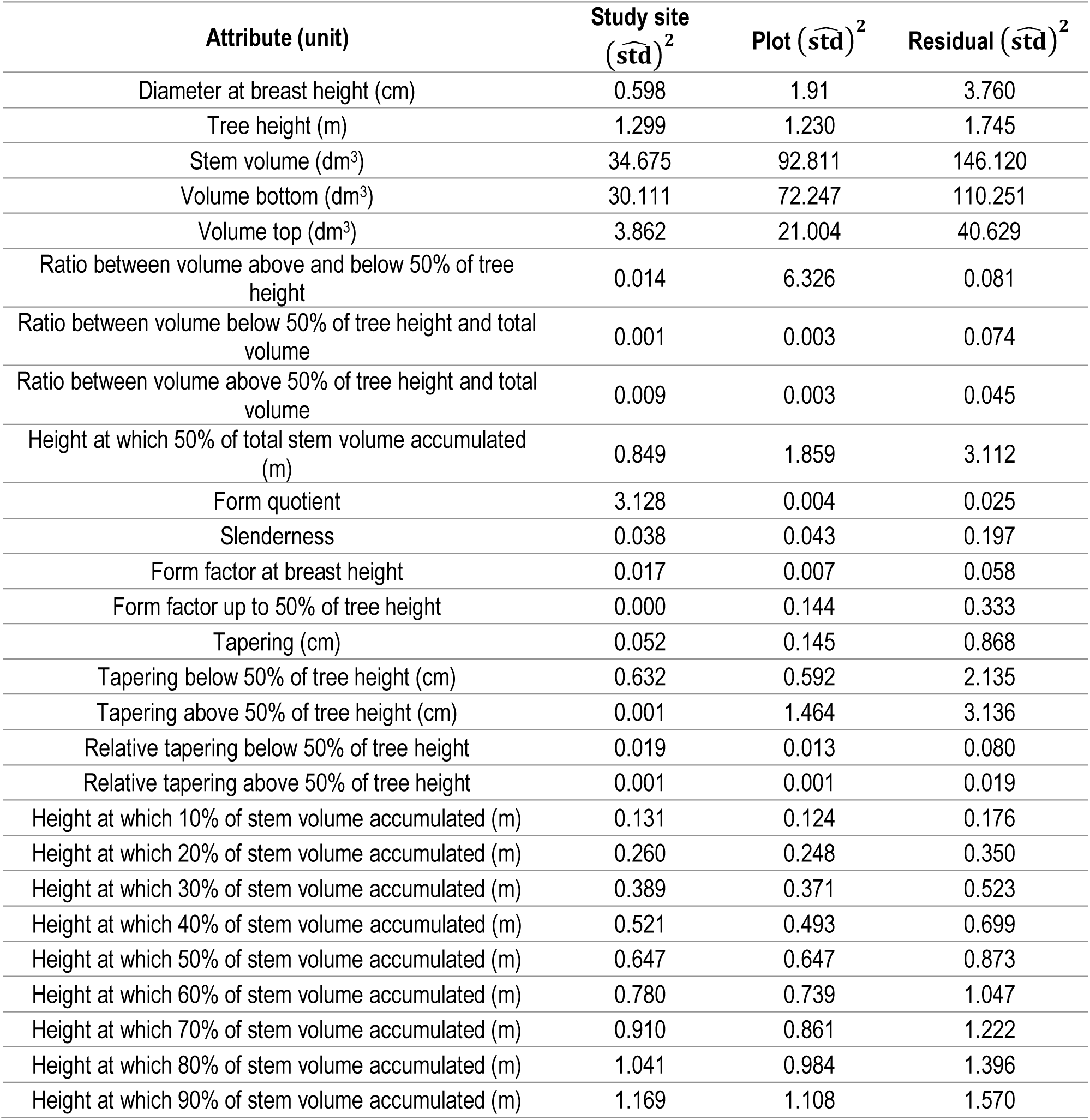
Variances obtained from the random part of the linear mixed-effects models at study site and plot level. Std = standard deviation.

| Attribute (unit) | Study site<br>($\widehat{\text{std}}$)<sup>2</sup> | Plot ($\widehat{\text{std}}$)<sup>2</sup> | Residual ($\widehat{\text{std}}$)<sup>2</sup> |
| --- | --- | --- | --- |
| Diameter at breast height (cm) | 0.598 | 1.91 | 3.760 |
| Tree height (m) | 1.299 | 1.230 | 1.745 |
| Stem volume (dm<sup>3</sup>) | 34.675 | 92.811 | 146.120 |
| Volume bottom (dm<sup>3</sup>) | 30.111 | 72.247 | 110.251 |
| Volume top (dm<sup>3</sup>) | 3.862 | 21.004 | 40.629 |
| Ratio between volume above and below 50% of tree height | 0.014 | 6.326 | 0.081 |
| Ratio between volume below 50% of tree height and total volume | 0.001 | 0.003 | 0.074 |
| Ratio between volume above 50% of tree height and total volume | 0.009 | 0.003 | 0.045 |
| Height at which 50% of total stem volume accumulated (m) | 0.849 | 1.859 | 3.112 |
| Form quotient | 3.128 | 0.004 | 0.025 |
| Slenderness | 0.038 | 0.043 | 0.197 |
| Form factor at breast height | 0.017 | 0.007 | 0.058 |
| Form factor up to 50% of tree height | 0.000 | 0.144 | 0.333 |
| Tapering (cm) | 0.052 | 0.145 | 0.868 |
| Tapering below 50% of tree height (cm) | 0.632 | 0.592 | 2.135 |
| Tapering above 50% of tree height (cm) | 0.001 | 1.464 | 3.136 |
| Relative tapering below 50% of tree height | 0.019 | 0.013 | 0.080 |
| Relative tapering above 50% of tree height | 0.001 | 0.001 | 0.019 |
| Height at which 10% of stem volume accumulated (m) | 0.131 | 0.124 | 0.176 |
| Height at which 20% of stem volume accumulated (m) | 0.260 | 0.248 | 0.350 |
| Height at which 30% of stem volume accumulated (m) | 0.389 | 0.371 | 0.523 |
| Height at which 40% of stem volume accumulated (m) | 0.521 | 0.493 | 0.699 |
| Height at which 50% of stem volume accumulated (m) | 0.647 | 0.647 | 0.873 |
| Height at which 60% of stem volume accumulated (m) | 0.780 | 0.739 | 1.047 |
| Height at which 70% of stem volume accumulated (m) | 0.910 | 0.861 | 1.222 |
| Height at which 80% of stem volume accumulated (m) | 1.041 | 0.984 | 1.396 |
| Height at which 90% of stem volume accumulated (m) | 1.169 | 1.108 | 1.570 |

**Table S3.** Statistically significant difference (p-value ≤ 0.05) caused by different thinning treatments and the following growth effect on the stem attributes characterizing post-thinning growth based on Tukey’s honest significance test. DBH = diameter at breast height, vol = total stem volume, vol_bottom = volume below 50% of tree height, vol_top = volume above 50% of tree height, r_vol = relative volume, r_vol_top = relative volume above 50% of tree height, h_vol50 = height at which 50% of stem volume accumulated, slend = slenderness, ff50 = form factor up to 50% of tree height, taper = tapering, taper_bottom = tapering below 50% stem height, taper_top = tapering above 50% of tree height. Please refer to Table 3 for descriptions of the attributes.

| Compared stem attributes caused by the thinning treatments and its growth response | | Attributes with statistically significant difference ($p\text{-value} \leq 0.05$) |
| --- | --- | --- |
| Moderate below | Moderate above | slend |
| Moderate below | Moderate systematic | slend |
| Moderate below | Intensive below | DBH, vol, vol_bottom, slend, taper, taper_bottom |
| Moderate below | Intensive above |  |
| Moderate below | Intensive systematic |  |
| Moderate below | No treatment | slend, taper_top, ff50 |
| Moderate above | Moderate systematic |  |
| Moderate above | Intensive below | DBH, vol, vol_bottom, r_vol, r_vol_top, h_vol50, slend, ff50, taper, taper_bottom, taper_top |
| Moderate above | Intensive above | slend |
| Moderate above | Intensive systematic | slend |
| Moderate above | No treatment |  |
| Moderate systematic | Intensive below | DBH, vol, vol_bottom, vol_top, h_vol50, slend, ff50, taper, taper_bottom, taper_top |
| Moderate systematic | Intensive above | slend |
| Moderate systematic | Intensive systematic | slend |
| Moderate systematic | No treatment |  |
| Intensive below | Intensive above | DBH, vol, vol_bottom, h_vol50, ff50, taper_bottom |
| Intensive below | Intensive systematic | DBH, vol, vol_bottom, r_vol, r_vol_top, h_vol50, slend, ff50, taper, taper_bottom, taper_top |
| Intensive below | No treatment | DBH, vol, vol_bottom, vol_top, h_vol50, slend, ff50, taper_bottom, taper_top |
| Intensive above | Intensive systematic |  |
| Intensive above | No treatment | slend |
| Intensive systematic | No treatment | slend |

## Notes

### Competing Interest Statement

The authors have declared no competing interest.

### Summary of Updates

- Methods revised - Table 1 revised - Supplementary materials added after the References

## References

1. Alonzo, M., Andersen, H.-E., Morton, D.C., Cook, B.D. 2018. Quantifying boreal forest structure and composition using UAV structure from motion. Forests 9(3): 119. https://doi.org/10.3390/f9030119

2. Aschoff, T., Thies, M., & Spiecker, H. 2004. Describing forest stands using terrestrial laser-scanning. International Archives of Photogrammetry, Remote Sensing and Spatial Information Sciences 35(5): 237–241.

3. Barbeito, I., Dassot, M., Bayer, D., Collet, C., Drössler, L., Löf, M., del Rio, M., Ruiz-Peinado, R., Forrester, D.I., Bravo-Oviedo, A., Pretzsch, H. 2017. Terrestrial laser scanning reveals difference in crown structure of *Fagus sylvatica* in mixed *vs*. pure European forests. Forest Ecology and Management 405: 381–390. https://doi.org/10.1016/j.foreco.2013.08.014

4. Bayer, D., Seifert, S., Pretsch, H. 2013. Structural crown properties of Norway spruce (*Picea abies* [L.] Karts.) and European beech (*Fagus sylvatica* [L.]) in mixed versus pure stands revealed by terrestrial laser scanning. Trees 27: 1035–1047. https://doi.org/10.1007/s00468-013-0854-4

5. Cabo, C., Ordóñez, C., López-Sánchez, C. A., Armesto, J. 2018. Automatic dendrometry: Tree detection, tree height and diameter estimation using terrestrial laser scanning. International journal of applied earth observation and geoinformation 69: 164–174. https://doi.org/10.1016/j.jag.2018.01.011

6. Calders, K., Newnhamn, G., Burt, A., Murpy, S., Raumonen, P., Herold, M., Culvenor, D., Avitabile, V., Disney, M., Armston, J., Kaasalainen, M. 2015. Nondestructive estimates of above-ground biomass using terrestrial laser scanning. Methods of Ecology and Evolution 6:198–208. https://doi.org/10.1111/2041-210X.12301

7. del Rio, M., Bravo-Oviedo, A., Pretzsch, H., Löf, M., Ruíz-Peinado, R. 2017. A review of thinning effects on Scots pine stands: From growth and yield to new challenges under global change. Forest Systems 28(2): eR03S. https://doi.org/10.5424/fs/2017262-11325

8. Eriksson, E. 2006. Thinning operations and their impact on biomass production in stands of Norway spruce and Scots pine. Biomass and Bioenergy 30: 848–854. https://doi.org/10.1016/j.biombioe.2006.04.001

9. Eriksson, H., Karlsson, K. 1997. Olika gallringsoch gödslingsregimers effecter på beståndsutvecklingen baserat på långliggande experiment i tall-och granbestånd i Sverige. Sveriges lantbruksuniversitet, Intsitutionen för skogsproduktion. Report No. 42: 1–135.

10. Farrar, J.L. 1961. Longitudinal variation in the thickness of the annual ring. Forestry Chronicle 37(4): 323–349. https://doi.org/10.5558/tfc37323-4

11. Fernández-Sarría, A., Velázquez-Martí, B., Sajdak, M., Martínez, L., Estornell, J. 2013. Residual biomass calculation from individual tree architecture using terrestrial laser scanned and ground-level measurements. Computers and Electronics in Agriculture 93: 90–97. https://doi.org/10.1016/j.compag.2013.01.012

12. Ferrarese, J., Affleck, D., Seielsatd, C. 2015. Conifer crown profile models from terrestrial laser scanning. Silva Fennica 49(1): 1106. https://doi.org/10.14214/sf.1106

13. Georgi, L., Kunz, M., Fichtner, A., Härdtle, W., Reich, K.F., Sturm, K., Welle, T., von Oheimb, G. 2018. Long-term abandonment of forest management has a strong impact on tree morphology and wood volume allocation pattern of European beech (*Fagus sylvatica* L.). Forests 9: 704. https://doi.org/10.3390/f9110704

14. Gonzalez de Tanago, J., Lau, A., Bartholomeus, H., Herol, M., Avitablile, V., Raumonen, P., Martius, C., Goodman, R.C., Disney, M., Manuri, S., Burt, A., Calders, K. 2017. Estimation of above-ground biomass of large tropical tree with terrestrial LiDAR. Methods in Ecology and Evolution 9(2): 223–234. https://doi.org/10.1111/2041-210X.12904

15. Goodbody, T.R., Coops, N.C., Marshall, P.L., Tompalski, P., Crawford, P. 2017. Unmanned aerial systems for precision forest inventory purposes: A review and case study. The Forestry chronicle 93(1): 71–81. https://doi.org/10.5558/tfc2017-012

16. Hackenberg, J., Morhart, C., Sheppard, J., Spiecker, H., Disney, M. 2014. Highly accurate tree models derived from terrestrial laser scan data: A method description. Forests 5(5):1069–1105. https://doi.org/10.3390/f5051069

17. Harper, J.L. 1977. Population biology of plants. Academic Press. London. 892 p.

18. Hauglin, M., Astrup, R., Gobakken, T., Næsset, E. 2013. Estimating single-tree branch biomass of Norway spruce with terrestrial laser scanning using voxel-based and crown dimension features. Scandinavian Journal of Forest Research 28(5): 456–469. https://doi.org/10.1080/02827581.2013.777772

19. Heinzel, J., Huber, M.O. 2017. Detecting tree stems from volumetric TLS data in forest environments with rich understory. Remote Sensing 9(1): 9. https://doi.org/10.3390/rs9010009 Jacobs, M., Rais, A., Pretzsch, H. 2020. Analysis of stand density effects on the stem form of Norway spruce trees and volume miscalculation by traditional form factor equations using terrestrial laser scanning (TLS). Canadian Journal of Forest Research 50: 51–64. https://doi.org/10.1139/cjfr-2019-0121

20. Isenburg, M. 2019. LAStools—Efficient LiDAR Processing Software, (version 181001 academic); rapidlasso GmbH: Gilching, Germany. Available online: http://rapidlasso.com/LAStools [accessed on December 19, 2019].

21. Juchheim, J., Ammer, C., Schall, P., Seidel, D. 2017a, Canopy space filling rather than conventional measures of structural diversity explains productivity of beech stands. Forest Ecology and Management 395: 19–26. https://doi.org/10.1016/j.foreco.2017.03.036

22. Juchheim, J., Annighöfer, P., Ammer, C., Calders, K., Raumonen, P., Seidel, D. 2017b. How management intensity and neighborhood composition affect the structure of beech (*Fagus sylvatica* L.) trees. Trees 31(5): 1723–1735. https://doi.org/10.1007/s00468-017-1581-z

23. Juodvalkis, A., Kairiukstis, L., Vasiliauskas, R. 2005. Effects of thinning on growth of six tree species in north-temperate forest of Lithuania. European Journal of Forest Research 124: 187–192. http://doi.org/10.1007/s10342-005-0070-x

24. Kankare, V., Holopainen, M., Vastaranta, M., Puttonen, E., Yu, X., Hyyppä, J., Vaaja, M., Hyyppä, H., Alho, P. 2013. Individual tree biomass estimation using terrestrial laser scanning. ISPRS Journal of Photogrammetry and Remote Sensing 75: 64–75. https://doi.org/10.1016/j.isprsjprs.2012.10.003

25. Kotivuori, E., Kukkonen, M., Mehtätalo, L., Maltamo, M., Korhonen, L., Packalen, P. 2020. Forest inventorie for small areas using drone imagery without in-situ field measurements. Remote Sensing of Environment 237: 111404. https://doi.org/10.1016/j.rse.2019.111404

26. Kozlowski, T. T. 1971. Growth and Development of Trees. Volume 2. Cambial Growth, Root Growth, and Reproductive Growth. Academic Press, New York, USA. 514 pp.

27. Kukkola, M., Mälkönen, E. 1997. The role of logging residues in site productivity after first thinning of Scots pine and Norway spruce stands. In: Hakkila, P., Heino, M., Puranen (eds.) Forest management for bioenergy. IEA Bioenergy, Proceedings of a joint meeting of Activities 1.1, 1.2 and 4.2 of Task XII in Jyväskylä, Finland, September 9 and 10, 1996. Metsäntutkimuslaitoksen tiedonantoja 640: 230–237.

28. Liang, X., Litkey, P., Hyyppä, J., Kaartinen, H., Vastaranta, M., Holopainen, M. 2011. Automatic stem mapping using single-scan terrestrial laser scanning. IEEE Transactions on Geoscience and Remote Sensing 50(2): 661–670. https://doi.org/10.1109/TGRS.2011.2161613

29. Liang, X., Kankare, V., Yu, X., Hyyppä, J., Holopainen, M. 2014. Automated stem curve measurement using terrestrial laser scanning. IEEE Transactions on Geoscience and Remote Sensing 52(3): 1739–1748. https://doi.org/10.1109/TGRS.2013.2253783

30. Liang, X., Hyyppä, J., Kaartinen, H., Lehtomäki, M., Pyörälä, J., Pfeifer, N., …, Wang, Y. 2018. International benchmarking of terrestrial laser scanning approaches for forest inventories. ISPRS Journal of Photogrammetry and Remote Sensing 144: 137–179. https://doi.org/10.1016/j.isprsjprs.2018.06.021

31. Luoma, V., Saarinen, N., Kankare, V., Tanhuanpää, T., Kaartinen, H., Kukko, A., Holopainen, M., Hyyppä, J., Vastaranta, M. 2019. Examining change in stem taper and volume growth with two-date 3D point clouds. Forests 10(5): 382. https://doi.org/10.3390/f10050382

32. Maas, H. G., Bienert, A., Scheller, S., Keane, E. 2008. Automatic forest inventory parameter determination from terrestrial laser scanner data. International Journal of Remote Rensing 29(5): 1579–1593. https://doi.org/10.1080/01431160701736406

33. Mäkinen, H., Isomäki, A. 2004a. Thinning intensity and growth of Scots pine stands in Finland. Forest Ecology and Management 201(2-3): 311–325. https://doi.org/10.1016/j.foreco.2004.07.016

34. Mäkinen, H., Isomäki, A. 2004b. Thinning intensity and long-term changes in increment and stem form of Norway spruce trees. Forest Ecology and Management 201(2-3): 295–309. https://doi.org/10.1016/j.foreco.2004.07.017

35. Mäkinen, H., Isomäki, A. 2004c. Thinning intensity and long-term changes in increment and stem form of Scots pine trees. Forest Ecology and Management 201(1-3): 21–34. https://doi.org/10.1016/j.foreco.2004.07.028

36. Metz, J., Seidel, D., Schall, P., Scheffer, D., Schulze, E.-D. 2013. Crown modelling by terrestrial laser scanning as an approach to assess the effect of aboveground intra- and interspecific competition on tree growth. Forest Ecology and Management 310: 275–288. https://doi.org/10.1016/j.foreco.2013.08.014

37. Meyer, F., Beucher, S. 1990. Morphological segmentation. Journal of visual communication and image representation 1(1): 21–46.

38. Mielikäinen, K., Valkonen, S. 1991. Harvennustavan vaikutus varttuneen metsikön tuotokseen ja tuottoihin Etelä-Suomessa. Summary: Effect of thinning method on the yield of middle-aged stands in southern Finland. In Finnish with English abstract. Folia Forestalia 776:1–22.

39. Montero, G., Cañellas, I., Ortega, C., Del Rio, M. 2001. Results from a thinning experiment in a Scots pine (*Pinus sylvestris* L.) natural regeneration stand in the Sistema Ibérico Mountain Range (Spain). Forest Ecology and Management 145: 151–161. https://doi.org/10.1016/S0378-1127(00)00582-X

40. National Land Survey of Finland. Finnref GNSS RINEX Service. 2020. Available online: https://www.maanmittauslaitos.fi/en/maps-and-spatial-data/positioning-services/rinex-palvelu [accessed January 7, 2020].

41. Nilsson, U., Agestam, E., Ekö, P-M., Elfving, B., Fahlvik, N., Johansson, U., Karlsson, K., Lundmark, T., Wallentin, C. 2010. Thinning of Scots pine and Norway spruce monocultures in Sweden–Effects of different thinning programmes on stand level gross- and net stem volume production. Studia Forestalia Suecia 219. 46 pp. ISSN 0039-3150, ISBN 978-91-86197-76-6. Noyer, E., Fournier, M., Constant, T., Collet, C., Dlouha, J. 2019. Biomechanical control of beech pole verticality (*Fagus sylvatica*) before and after thinning: theoretical modelling and ground-truth data using terrestrial LiDAR. American Journal of Botany 106(2): 187–198. https://doi.org/10.1002/ajb2.1228

42. Pinheiro, J., Bates, D., DebRoy, S., Sarkar, D., R Core Team. 2016. nlme: Linear and Nonlinear Mixed Effects Models. Available: http://CRAN.R-project.org/package=nlme [accessed January 21, 2020] R package version 3.1-143.

43. Popescu, S. C., Wynne, R. H. 2004. Seeing the trees in the forest. Photogrammetric Engineering & Remote Sensing 70(5): 589–604. https://doi.org/10.14358/PERS.70.5.589

44. Pukkala, T., von Gaudow, K. (Eds.) 2012. Continuous cover forestry. 2nd Edition. Springer Dordrecht Heidelberg London New York. 295 p.

45. Puliti, S., Ørka, H.O., Gobakken, T., Næsset, E. 2015. Inventory of small forest areas using an unmanned aerial system. Remote Sensing 7(8): 9632–9654. https://doi.org/10.3390/rs70809632

46. Pyörälä, J., Kankare, V., Liang, X., Saarinen, N., Rikala, J., Kivinen, V.-P., Sipi, M., Holopainen, M., Hyyppä, J., Vastaranta, M. 2019. Assessing log geometry and wood quality in standing timber using terrestrial laser-scanning point clouds. Forestry 92(2): 177–189. https://doi.org/10.1093/forestry/cpy044

47. R Core Team. 2019. R: A language and environment for statistical computing. Vienna, Austria: R Foundation for Statistical Computing. https://www.R-project.org [accessed January 21, 2020]

48. Rantala, S. (ed.) 2011. Finnish forestry practice and management. Metsäkustannus. Helsinki. 271 pp.

49. Raumonen, P., Kaasalainen, M., Åkerblom, M., Kaasalainen, S., Kaartinen, H., Vastaranta, M., Holopainen, M., Disney, M., Lewis, P. 2013. Fast automatic precision tree model from terrestrial laser scanning. Remote Sensing 5(2): 491–520. https://doi.org/10.3390/rs5020491

50. Saarinen, N., Kankare, V., Vastaranta, M., Luoma, V., Pyörälä, J., Tanhuanpää, T., Liang, X., Kaartinen, H., Kukko, A., Jaakkola, A., Yu, X., Holopainen, M., Hyyppä, J. 2017. Feasibility of terrestrial laser scanning for collecting stem volume information from single trees. ISPRS Journal of Photogrammetry and Remote Sensing 123:140–158. https://doi.org/10.1016/j.isprsjprs.2016.11.012

51. Saarinen, N., Vastaranta, M., Näsi, R., Rosnell, T., Hakala, T., Honkavaara, E., Wulder, M.A., Luoma, V., Tommaselli, A.M.G., Imai, N.N., Ribeiro, E.A.W., Guimarães, R.B., Holopainen, M., Hyyppä, J. 2018. Assessing biodiversity in boreal forests with UAV-based photogrammetric point clouds and hyperspectral imaging. Remote Sensing 10(2): 338. 338; https://doi.org/10.3390/rs10020338

52. Saarinen, N., Kankare, V., Pyörälä, J., Yrttimaa, T., Liang, X., Wulder, M.A., Holopainen, M., Hyyppä, J., Vastaranta, M. 2019. Assessing the effects of sample size on parametrizing a taper curve equation and the resultant stem-volume estimates. Forests 10(10): 848. https://doi.org/10.3390/f10100848

53. Savill, P.S., Sandels, A.J. 1983. The influence of early respacing on the wood density of Sitka spruce. Forestry 56(2): 109–120. https://doi.org/10.1093/forestry/56.2.109

54. Seidel D. 2018. A holistic approach to determine tree structural complexity based on laser scanning data and fractal analysis. Ecology and Evolution 8(1): 128–134. https://doi.org/10.1002/ece3.3661

55. Seidel, D., Ruzicka, K.J., Puettmann, K. 2016. Canopy gaps affect the shape of Douglas-fir crown in the western Cascades, Oregon. Forest Ecology and Management 363: 31–38. https://doi.org/10.1016/j.foreco.2015.12.024

56. Srinivasa, S., Popescu, S.C., Eriksson, M., Sheridan, R.D., Ku, N.-K. 2014. Multi-temporal terrestrial laser scanning for modelling tree biomass change. Forest Ecology and Management 318: 304–317. https://doi.org/10.1016/j.foreco.2014.01.038

57. Stiers, M., Willia, K., Seidel, D., Ehbrecht, M., Kabal, M., Ammer, C., Annighöfer, P. 2018. A quantitative comparison of the structural complexity of managed, lately managed and promari European beech (*Fagus sylvatica* L.) forests. Forest Ecology and Management 430: 357–365. https://doi.org/10.1016/j.foreco.2018.08.039

58. Stovall, A.E.L., Shugart, H.H. 2018. Improved biomass calibration and validation with terrestrial LiDAR: Implications of future LiDAR and SAR missions. IEEE Journal of Selected Topics in Applied Earth Observations and Remote Sensing 11(10): 3527–3537. https://doi.org/10.1109/JSTARS.2018.2803110

59. Valinger, E. 1992. Effects of thinning and nitrogen fertilization on stem growth and stem form of *Pinus Sylvestris* trees. Scandinavian Journal of Forest Research 7(1-4): 219–228. https://doi.org/10.1080/02827589209382714

60. Valinger, E., Sjögren, H., Nord, G., Cedergren, J. 2019. Effects of stem growth of Scots pine 33 years after thinning and/or fertilization in northern Sweden. Scandinavian Journal of Forest Research 31(1): 33–38. https://doi.org/10.1080/02827581.2018.1545920

61. Viljanen, N., Honkavaara, E., Näsi, R., Hakala, T., Niemeläinen, O., Kaivosoja, J. 2018. A Novel Machine Learning Method for Estimating Biomass of Grass Swards Using a Photogrammetric Canopy Height Model, Images and Vegetation Indices Captured by a Drone. Agriculture 8: 70. https://doi.org/10.3390/agriculture8050070

62. Vuokila, Y. 1977. Selective thinning from above as a fact of growth and yield. Folia Forestalia 298: 1-26. (In Finnish with English summary)

63. Wallace, L., Lucieer, A., Malenovský, Z., Turner, D., Vopenka, P. Assessment of forest structure using two UAV techniques: A comparison of airborne laser scanning and structure from motion (SfM) point clouds. Forests 7(3): 62. https://doi.org/10.3390/f7030062

64. White, J. 1980. Demographic factors in population of plants. In: Solbrig, O.T. (Ed.) Demography and Evalutaion in Plant Populations. Botanical Monographs 15. University of California Press. Berkeley and Los Angeles, USA. 222 p.

65. Wilkes, P., Lau, A., Disney, M., Calders, K., Burt, A., de Tanago, J. G., …, Herold, M. 2017. Data acquisition considerations for terrestrial laser scanning of forest plots. Remote Sensing of Environment 196: 140–153. https://doi.org/10.1016/j.rse.2017.04.030

66. Yrttimaa, T., Saarinen, N., Kankare, V., Liang, X., Hyyppä, J., Holopainen, M., Vastaranta, M. 2019. Investigating the feasibility of multi-scan terrestrial laser scanning to characterize tree communities in southern boreal forests. Remote Sensing 11(12): 1423. https://doi.org/10.3390/rs11121423

67. Yrttimaa, T., Saarinen, N., Kankare, V., Hynynen, J., Huuskonen, S., Holopainen, M., Hyyppä, J., Vastaranta, M. 2020. Performance of terrestrial laser scanning to characterize managed Scots pine (Pinus sylvestris L.) stands is dependent on forest structural variation. EarthArXiv March 5. https://doi.org/10.31223/osf.io/ybs7c

68. Yrttimaa, T., Saarinen, N., Kankare, V., Viljanen, N., Hynynen, J., Huuskonen, S., Holopainen, M., Hyyppä, J., Honkavaara, E., Vastaranta, M. 2020. Multisensorial close-range sensing generates benefits for characterization of managed Scots pine (*Pinus sylvestris* L.) stands. ISPRS International Journal of Geo-Information 9(5): 309. https://doi.org/10.3390/ijgi9050309

69. Zhang, W., Wan, P., Wang, T., Cai, S., Chen, Y., Jin, X., & Yan, G. 2019. A novel approach for the detection of standing tree stems from plot-level terrestrial laser scanning data. Remote sensing 11(2): 211. https://doi.org/10.3390/rs11020211

